# RegFM: an interpretable context-aware foundation model for human transcriptional regulation

**DOI:** 10.64898/2026.08.17.744355

**Authors:** Zijing Gao, Yining Sun, Haochen Wang, Rui Jiang, Qiao Liu

## Abstract

Transcriptional regulation is governed by interactions between cis-regulatory elements (CREs) and trans-acting regulators in a context-specific manner. Although DNA and single-cell foundation models have enabled modeling regulatory biology at scale, most represent either sequence or cellular state alone, limiting their ability to capture context-dependent gene regulation. Here we present RegFM, a context-aware foundation model for human transcriptional regulation. RegFM treats transcriptional regulation as a *dialogue* between cis-regulatory sequences (e.g., CREs) and trans-acting regulators (e.g., transcription factors (TFs) and chromatin regulators (CRs)) by coupling long-range CRE representations with TFs and CRs activity. Trained on large-scale ENCODE and CELLxGENE transcriptomic profiles, RegFM learns gene-centered regulatory representations that generalize across unseen cellular contexts. In a wide range of tasks, including gene expression prediction, cis-regulatory element annotation, bivalent promoter and dosage-sensitivity classification, and perturbation-response prediction, RegFM consistently improves over existing methods. RegFM emerges as a scalable and interpretable framework for modeling human transcriptional regulation and provides insights into context-dependent gene regulatory programs.

## Introduction

Transcriptional regulation links genomic sequence to cell-type-specific gene expression programs and is shaped by interactions among cis-regulatory elements (CREs), transcription factors (TFs) and chromatin regulators (CRs). Many disease-associated variants mainly reside in non-coding regions and influence disease risk by disrupting regulatory programs rather than altering protein sequence^1–4^. A central challenge is that gene regulation is highly context dependent: the same DNA sequence can be interpreted differently across tissues, cell types^5^. Therefore, mechanistic modeling of gene regulation needs to integrate both cis-regulatory sequence and the trans-regulatory cellular environment. Large-scale datasets, including ENCODE^6^ and CELLxGENE^7^, provide an unprecedented opportunity to learn such context-dependent regulatory mechanism at scale^8^.

Task-specific computational models have addressed important aspects of transcriptional regulation, including gene expression prediction and gene regulation networks (GRNs) inference^9, 10^. However, these approaches are typically optimized for specific tasks, limiting the applications in broader biological contexts. The rapid development of genomic foundation models has reshaped computational genomics by learning transferable representations from large-scale genomic data^11^. Two complementary directions have emerged. DNA foundation models, such as DNABERT^12, 13^, the Nucleotide Transformer (NT)^14^, EVO^15, 16^, Enformer^17^, and AlphaGenome^18^ learn representations directly from DNA sequence and can capture regulatory syntax and long-range sequence dependency. However, DNA sequence alone does not determine regulatory activity: the same CRE may be active, inactive, or exert different effects depending on the cellular context. Conversely, single-cell foundation models, including scFoundation^19^, Geneformer^20^, scGPT^21^, and EpiAgent^22^ learn cellular representations from transcriptomic or epigenomic single-cell atlas data. These models are powerful for characterizing cellular state, but they largely neglect the DNA sequences that encode the cis-regulatory potential. Therefore, current genomic foundation models typically represent either the DNA sequence or the cellular state, but not their context-dependent regulatory interaction.

Toward this goal, our prior work EpiGePT^23^ and the recent work Corgi^24^ have demonstrated the effectiveness of integrating DNA sequence with trans-regulatory profiles for predicting genome-wide epigenomic tracks. GET^5^ represents the closest foundation-scale effort toward linking regulatory context with transcriptional activity by integrating chromatin accessibility and gene expression across human cell types. However, GET remains constrained by how it defines and represents regulatory context. First, GET requires paired ATAC-seq and RNA-seq data as input, which are substantially less available than RNA-seq alone, limiting the scale and diversity of applicable cellular contexts. Second, its trans-regulatory component is primarily represented through motif features within accessible regions, which makes it difficult to resolve homologous TFs with similar motifs but distinct expression patterns. These limitations highlight the need for a foundation model that directly couples long-range cis-regulatory sequence with trans-regulatory activities to enable context-aware modeling of transcriptional regulation.

To address these limitations, we develop RegFM, a context-aware genomic foundation model for human transcriptional regulation. RegFM models the transcriptional activity of each gene by pairing a long-range gene-centered DNA sequence with the activity profile of TFs and CRs, and integrates these modalities through a cross-attention module to learn context-dependent cis-trans regulatory interactions. Using self-supervised language modeling and supervised expression prediction on large-scale ENCODE and CELLxGENE transcriptomic datasets, RegFM learns gene-centered regulatory representations across diverse human cell types and tissues. Across gene expression prediction, CRE annotation, bivalent promoter and gene dosage sensitivity prediction, and genetic perturbation response prediction tasks, RegFM consistently outperforms existing genomic foundation models and task-specific baselines. Beyond prediction, the learned cross-attention maps reveal regulatory relationships among regulatory sequence, trans-acting regulators and target genes, providing an interpretable view of context-specific transcriptional landscape. RegFM represents a scalable and interpretable framework for modeling human transcriptional regulation and provides insights into the cis-trans regulatory mechanisms that shape context-specific gene expression.

### Results Overview

RegFM is a context-aware genomic foundation model that jointly encodes DNA sequence and cellular context to model transcriptional regulation. To learn generalizable regulatory rules across diverse cellular states, RegFM was pretrained on two complementary resources, including 1,018 bulk RNA-seq experiments from the ENCODE consortium and pseudo-bulk RNA-seq profiles generated from 30 million cells spanning 663 cell types from CellxGene (**Fig. 1a** and section “Methods”). Together, these resources provide RegFM with rich and diverse information for learning general regulatory patterns across different cell types. RegFM adopts a dual-transformer architecture with a *cis*-DNA transformer module and a *trans*-context transformer module coupled by a cis-trans cross-attention module. The *cis*-DNA transformer processes a gene-center long DNA sequence as input where the sequence is tokenized into non-overlapping k-mers using a BPE tokenizer (Fig. 1b). The *trans*-context transformer consists of 2,103 selected TFs and CRs curated from Cis-BP^25^ and CRdb^26^. This module is designed to summarize the trans-acting regulatory context environment that determines which CREs are activated.

**Fig. 1.**
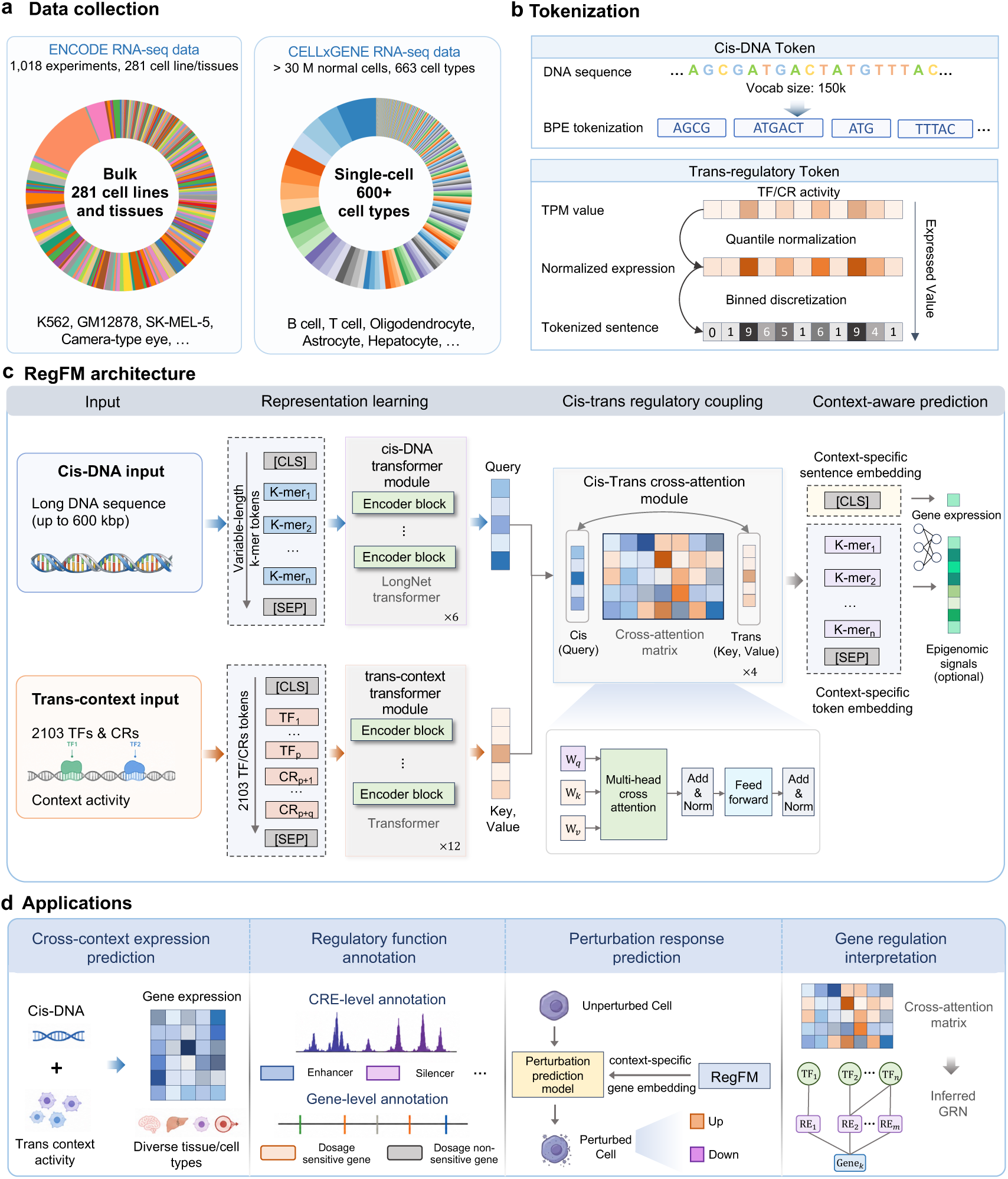
Overview of RegFM. **a**, Data collection for training RegFM, consisting of 1,018 RNA-seq experiments from ENCODE consortium, covering 281 cell lines or tissues, together with pseudo-bulk RNA-seq data aggregated from about 30 million normal cells from CellxGene, spanning 663 cell types. **b**, Tokenization strategy for cis-DNA transformer module and trans-context transformer module. DNA sequences are tokenized using BPE tokenizer, and TF/CR expression values are processed through logarithmic transformation, quantile normalization, and binned discretization. **c**, The model architecture of RegFM. For gene expression prediction task, a cis-trans cross-attention module is designed to integrate the cis-DNA and trans-context information. A multi-layer perceptron (MLP) is subsequently applied to the classification token (‘CLS’) embedding to predict cell-type-specific gene expression values. **d,** The primary application scenarios of RegFM, including cross-context gene expression prediction, regulatory function annotation, genetic perturbation transcriptional response prediction, and gene regulatory network inference.

To model the context-specific transcriptional activity, RegFM introduces the cis-trans cross-attention module to explicitly model the coupling between CREs and trans-regulatory factors (Fig. 1c). Conceptually, the cis-DNA transformer module encodes the regulatory grammar of gene-centered DNA sequences, defining which CREs could potentially be engaged. The trans-context transformer encodes the cellular regulatory state, specifying which TFs and CRs are available and active in that context. Through cis-trans cross-attention module, RegFM learns a context-specific *dialogue* between cis-regulatory elements and trans-acting regulators, dynamically prioritizing which CREs are likely to contribute to transcriptional output under a given trans-regulatory environment. Importantly, the learned cross-attention maps provide interpretable regulatory relationships between CREs and TFs/CRs, offering biologically meaningful interpretation about context-specific regulatory mechanisms.

### RegFM accurately predicts gene expression in unseen cellular contexts

To comprehensively benchmark the performance of RegFM in predicting gene expression in unseen cellular contexts, we first partitioned the cell type-by-gene matrix into training and test sets by randomly selecting 20% of cellular contexts as the test set. (**Fig. 2a**). We compared RegFM with DNA foundation models, including DNABERT^12^, DNABERT-2^13^, the Nucleic Transformer^14^, the Nucleotide Transformer^27^, EVO-2^16^, and a task-specific gene expression prediction model, Xpresso^10^. To ensure a fair comparison, all models were fine-tuned or trained using the same training data (Supplementary Text S1). The primary evaluation metrics included the Pearson correlation coefficient (PCC), Spearman correlation coefficient (SCC), *R*^2^, mean absolute error (MAE), and root mean squared error (RMSE). We conducted both context-wise and gene-wise evaluations (see the *Methods* section).

**Fig. 2.**
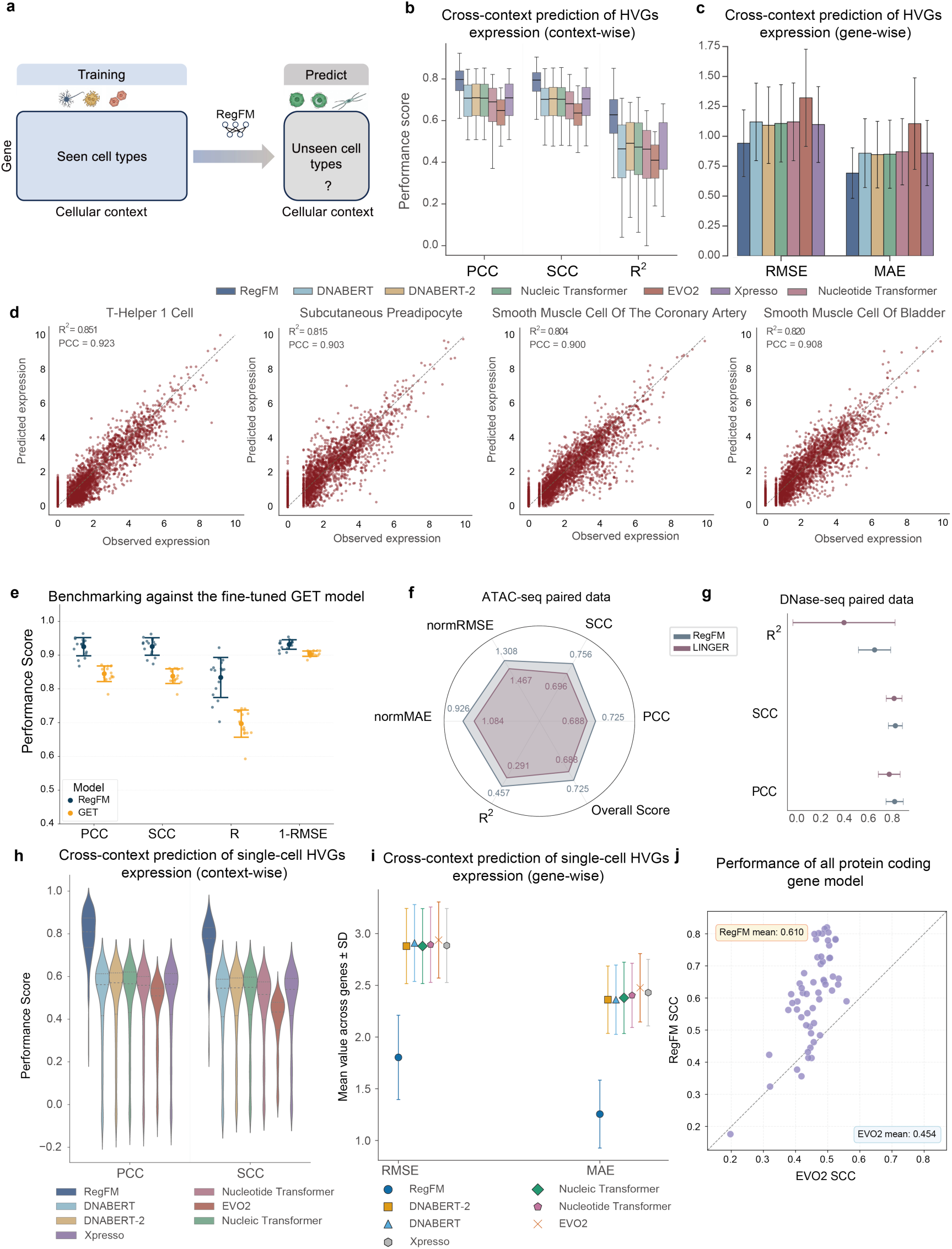
Benchmarking RegFM for gene expression prediction compared with baseline methods. **a,** Schematic illustration of the data partition strategy. **b,** Cross-cellular context prediction performance for highly variable genes (n = 3,000) on bulk RNA-seq data from ENCODE, evaluated by PCC, SCC, and *R*^2^. **c,** Gene-wise prediction performance evaluated by RMSE and MAE. **d,** Scatter plots showing the distribution of predicted versus observed expression values for RegFM across four representative cell lines. **e,** Comparison of gene expression prediction performance between RegFM and GET on 16 unseen cell types in the PBMC 10x dataset. **f,** Comparison of HVG (n = 1,000) expression prediction performance between RegFM and LINGER on 26 cell line/tissues with paired ATAC-seq and RNA-seq data. **g,** Comparison of prediction performance between RegFM and LINGER on 111 cell lines and tissues with paired DNase-seq and RNA-seq data. **h,** Cross-cellular context prediction performance on pseudo-bulk RNA-seq data from CELLxGENE, evaluated by PCC, SCC, and *R*^2^. **i,** Gene-wise prediction performance on the CELLxGENE dataset evaluated by RMSE and MAE. **j,** Comparison of prediction performance between RegFM and EVO-2 across 56 test cell lines using all protein-coding genes, evaluated by SCC.

As shown in **Fig. 2b-c**, when predicting the expression of 3,000 highly variable genes (HVGs) in 56 unseen cellular contexts, RegFM significantly outperforms the baseline methods (one-sided paired Wilcoxon signed-rank tests *p*-value < 4.30 × 10^−9^) in both context-wise and gene-wise evaluations. For example, RegFM achieved an average PCC of 0.774, substantially outperforming the second-best method (DNABERT-2) with a PCC of 0.695. In unseen cell type, such as T helper-1, RegFM achieved a PCC of 0.923 and an *R*^2^ of 0.851 (**Fig. 2d**).

Since GET^5^ requires paired chromatin-accessibility and transcriptomic profiles, it could not be included in the ENCODE benchmark where matched accessibility data were not consistently available. Therefore, we compared RegFM to GET in gene expression prediction by training RegFM on the same PBMC 10x dataset under a leave-one-out setting across 16 cell types used by GET. Despite the fact that GET additionally incorporates scATAC-seq peak information, RegFM still achieved better predictive performance with the average PCC improvement from 0.845 to 0.925 (**Fig. 2e**). In the held-out CD4 Tcm cell type, RegFM achieved a PCC of 0.968, substantially exceeding the GET model fine-tuned on the same dataset (0.740) (Supplementary Figs. S7a-b). These comparisons indicate that RegFM infers context-specific transcriptional activity from gene-centered sequence and TF/CR activity profiles without requiring additional chromatin-accessibility input.

In addition, we compared RegFM with LINGER, a task-specific multi-omics regulatory model that uses paired gene-expression and chromatin-accessibility data to model target gene expression and infer GRNs. We curated 26 cellular contexts with matched ATAC-seq and RNA-seq data, as well as 111 cellular contexts with matched DNase-seq and RNA-seq data from ENCODE. For paired ATAC-seq paired datasets, RegFM consistently outperformed LINGER across evaluation metrics, increasing the overall score from 0.696 to 0.756 (**Fig. 2f**, see Methods). Similarly, on the DNase-seq paired dataset, we conducted five-fold cross-validation on cellular contexts, and RegFM consistently outperforms LINGER, achieving average PCC of 0.843, compared to 0.797 of LINGER (**Fig. 2g**).

We further train a RegFM model on large-scale scRNA-seq atlas data from CELLxGENE using scRNA-seq profiles of ∼43 million cells (∼30 million normal cells). Single cells were aggregated into pseudo-bulk profiles comprising 663 cell types (see the *Methods* section). Across 133 unseen cell types and 1,000 HVGs, RegFM substantially outperformed baseline methods by a large margin in both context-wise and gene-wise evaluations (**Fig. 2h-i**). For example, RegFM achieves an average PCC of 0.789 in unseen cell types, remarkedly higher than the second-best method with a PCC of 0.507 (one-sided paired Wilcoxon signed-rank tests *p*-value = 2.83 × 10^−2^^3^). Finally, we trained a full-scale model on ENCODE covering all protein-coding genes, comprising 3.6 million training instances, to support downstream applications and analyses. This model continued to outperform the DNA foundation model EVO-2, achieving an average SCC of 0.610 compared to 0.454 of EVO-2 (**Fig. 2j**). The results demonstrate that RegFM scales effectively to genome-wide expression modeling and learns transferable regulatory representations across diverse human cell types.

### Context-aware RegFM embeddings facilitate diverse downstream genomics tasks

Beyond gene-expression prediction, we asked whether RegFM learns regulatory embeddings that could benefit to broader functional genomics tasks. We used RegFM-derived embeddings of CREs and target genes as inputs to task-specific prediction heads and evaluated the performance across diverse functional genomics tasks, including CRE identification, cell-type-specific classification of enhancers, silencers and insulators, gene dosage sensitivity prediction, and promoter bivalency prediction (**Fig. 3a**).

**Fig. 3.**
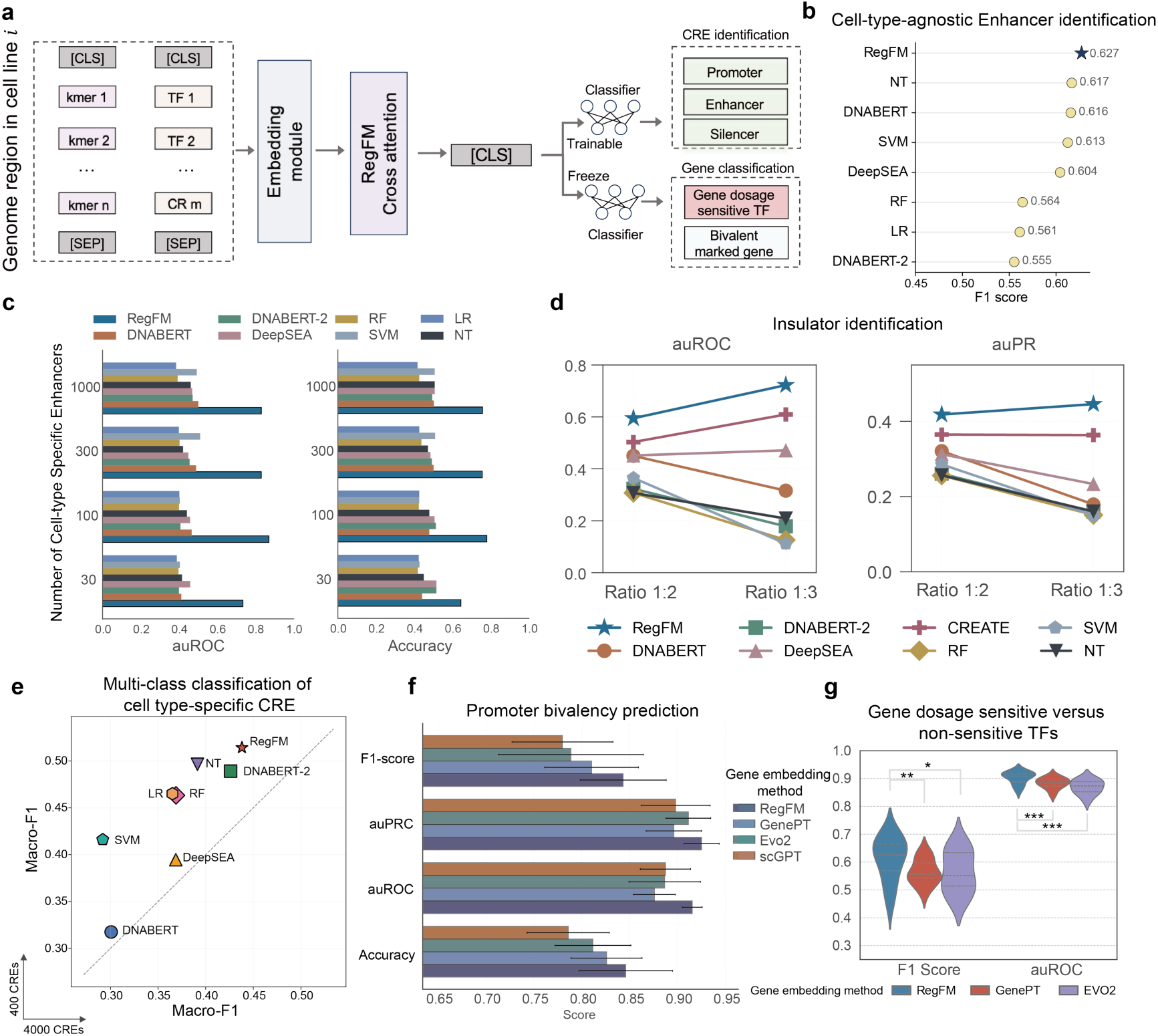
Fine-tuning RegFM with cell-type-specific genomic region embedding. **a.** Workflow for downstream genomic functional annotation tasks using fine-tuned RegFM (enhancer identification, gene dosage sensitivity prediction, and promoter bivalency prediction). **b,** F1 score for enhancer prediction performance across 44 cell lines and tissues in a cell-type-agnostic setting of RegFM and baseline methods, distinguishing EnhancerAtlas annotations from non-enhancer accessible chromatin regions. **c,** auROC and accuracy for identifying fully cell-type-specific enhancers across 44 cell lines and tissues using the same set of CREs and background regions, evaluated for fine-tuned RegFM and baseline methods across varying numbers of CREs (30, 100, 300, and 1000). **d.** The performance (auROC and auPR) of RegFM and baseline methods in insulator identification across the GM12878, Hela-S3, HepG2, and K562 cell lines. **e,** The performance (macro-averaged F1 score) of RegFM and baseline methods on cell-type-specific multi-class classification of regulatory elements, including enhancers, silencers, insulators, and background regions. **f,** Performance of fine-tuned RegFM and baseline methods in distinguishing bivalent versus non-bivalent genes from a previous study^30^. **g,** Performance (F1-score, auPRC, auROC, Accuracy) of RegFM fine-tuned to distinguish dosage-sensitive versus dosage-insensitive TFs, compared to alternative methods that provide gene embedding (GenePT, EVO2).

### Cell-type-agnostic and cell-type-specific CRE identification

Enhancer annotations for 44 cell lines/tissues were collected from EnhancerAtlas^28, 29^. As shown in **Fig. 3b**, the fine-tuned RegFM achieved an F1 score of 0.627, outperforming DNABERT (0.616), DeepSEA (0.345), and an SVM classifier (0.613) in cell-type-agnostic enhancer identification (see *Methods*). We next evaluated RegFM under a more challenging cell-type-specific setting by randomly sampling 8,938 CREs. Pairwise enhancer overlap across cell types ranged from 19% to 69%. For each cell type, negative examples were preferentially selected from CREs annotated as enhancers in multiple other cell types, thereby introducing conflicting activity labels across cellular contexts. RegFM again outperformed all baseline methods, achieving an auROC of 0.843 and an auPR of 0.860 (Supplementary Figs. S12a).

Furthermore, we considered a matched-region setting in which the same genomic regions were evaluated across multiple cell types but assigned context-dependent regulatory labels. As shown in **Fig. 3c**, across 44 cell types and different numbers of enhancers (30, 100, 300, 1,000), RegFM consistently outperformed baseline methods. For example, in the setting with 1,000 CREs, RegFM achieved an auROC of 0.831 while baselines performed near chance since their embeddings cannot distinguish cell-type-specific enhancer activity at the same genomic loci. Similarly, we collected insulator and background annotations from 4 cell lines, HepG2, K562, Hela-S3, and GM12878, from Cui et. al^4^. In the classification task for insulator, RegFM consistently outperformed baseline methods in handling different positive-to-negative sample ratios (1:2 and 1:3) (**Fig. 3d**). We also observed that CREATE, a dedicated CRE prediction model, outperforms other baseline methods, but remained lower than RegFM (Supplementary Text S2 and Fig. S11). These results demonstrate that RegFM captures both shared and cell-type-specific regulatory element activity, enabling context-aware annotation of the genome-wide CREs.

### Multi-class classification of cell type-specific CRE

To further evaluate the ability of RegFM to distinguish different classes of context-specific CREs, we performed multi-class classification of enhancers, silencers, insulators and background regions across four cell lines: HepG2, GM12878, HeLa-S3 and K562. CREs from the four cell lines were collected from Cui et al^4^ and randomly split into training and test sets. As shown in **Fig. 3e**, RegFM achieved the highest macro-averaged F1 score across different sample size settings. With 4000 samples per cell line, RegFM reached a macro-averaged F1 score of 0.438, outperforming the best-performing baseline (0.426).

### Bivalent marked gene classification

Bivalent promoters, characterized by the coexistence of activating and repressive histone modifications, represent a gene-level regulatory state associated with stem cell pluripotency^20, 30^. We therefore tested whether RegFM-derived gene embeddings could distinguish bivalently marked genes from genes with non-bivalent promoter states in embryonic stem cells (ESCs). Using the gene sets previously reported^30^ along with the matched gene expression data in ESCs (see *Methods*), RegFM significantly outperformed the baseline methods GenePT, EVO-2, and scGPT^21^, which generate gene embeddings from textual descriptions of gene function, DNA sequences, or gene expression profile, respectively. In five-fold cross-validation, RegFM achieved the highest average F1 score of 0.843, compared with 0.810 for GenePT and 0.789 for EVO-2 (**Fig. 3f**). These results suggest that RegFM embeddings capture gene-level regulatory features associated with promoter bivalency beyond sequence-or annotation-derived gene representations.

### Gene dosage sensitivity prediction

A major challenge in interpreting copy number variants (CNVs) during genetic diagnosis is determining which genes are sensitive to dosage alterations^20^. We therefore evaluated whether RegFM-derived gene embeddings could predict dosage-sensitive transcription factors. For each candidate gene, RegFM generated context-specific embeddings across 281 ENCODE cellular contexts, which were integrated using an attention-based fusion module and passed to a classification head. (see *Methods*) Using gene sets reported in previous studies^20, 31^, RegFM significantly outperformed all baseline methods in 10-fold cross-validation (**Fig. 3g**), achieving an average F1-score of 0.610 and an average auROC of 0.903. The second-best method, GenePT, achieved an average F1-score of 0.564 (one-sided paired t-test, *p*-value < 0.01) and auROC of 0.883 (one-sided paired t-test, *p*-value < 0.001). These results suggest that integrating regulatory representations across cellular contexts improves the identification of dosage-sensitive regulators, highlighting the value of RegFM for gene-level functional prioritization.

Together, these results demonstrate that RegFM-derived embeddings encode regulatory information that can be utilized in a wide range of functional genomic prediction tasks. By integrating cis-regulatory sequence with cell-type-specific regulator activity, RegFM captures both sequence-encoded potential and context-dependent regulatory activity, enabling accurate annotation of CREs, promoter bivalency and dosage-sensitive regulators across cellular contexts.

### Context-aware RegFM embeddings improve perturbation-response prediction

Predicting transcriptional responses to genetic perturbations requires representations that capture both the perturbed gene and the cellular context in which the perturbation occurs. Recent perturbation-response models have increasingly used gene embeddings derived from large language models^32^, biological knowledge graphs^33, 34^, and single-cell foundation model^21^ to represent perturbation targets. Although useful, these embeddings are typically static across cell types and therefore do not explicitly encode the regulatory environment that shapes perturbation outcomes. We therefore asked whether RegFM embeddings could improve perturbation-response prediction. To test this, we incorporated RegFM into the lightweight Scouter framework^32^ by integrating the static gene embedding from GenePT^35^ with RegFM-derived context-specific gene embedding 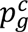 (**Fig. 4a**; see *Methods*; Supplementary Text S3). Four baseline methods were selected for comparison: GEARS^34^, biolord^33^, scGPT^21^, and original Scouter^32^, covering knowledge graph-based, transcriptome-based, and literature knowledge-based approaches.

To evaluate prediction of responses to unseen single-gene perturbations, we benchmarked RegFM and baseline methods on five processed Perturb-seq datasets from PerturBase^36^: Brain^37^, MCF10A^38^, Neurons^39^, embryonic stem cells^40^, and foreskin fibroblasts^41^. Following Scouter, we evaluated post-perturbation expression prediction using normalized mean squared error (normMSE) and SCC, and PCC of gene expression change over control cells (see *Methods* section). Across the five Perturb-seq datasets, evaluated under a 5-fold cross-validation setting over perturbation genes, RegFM consistently achieves the best performance. In the Brain dataset, for example, it achieved the average SCC of 0.955, outperforming biolord (0.931) and GEARS (0.904) (**Fig. 4b**). As shown in **Fig. 4c**, RegFM substantially outperforms all baseline methods in predicting transcriptional responses to unseen perturbations in the MCF10A cell line. When evaluating gene expression changes, RegFM achieves an average PCC of 0.789 on the top 20 differentially expressed genes (DEGs), whereas the best baseline, GEARS achieved 0.563. For non-zero expressed genes, RegFM also reaches a PCC of 0.443, compared to 0.251 for scGPT. Consistently, RegFM maintains superior performance across all baseline methods in predicting absolute post-perturbation gene expression levels (**Fig. 4d**). Furthermore, RegFM not only improves the accuracy of post-perturbation expression prediction but also better captures the direction of transcriptional changes. In hESCs following SMAD3 perturbation, RegFM correctly predicts the direction of change for the top 15 DEGs, including downregulated genes such as COA6, which original Scouter incorrectly predicts as upregulated (**Fig. 4e**, Supplementary Figs. S9a-b).

**Fig. 4.**
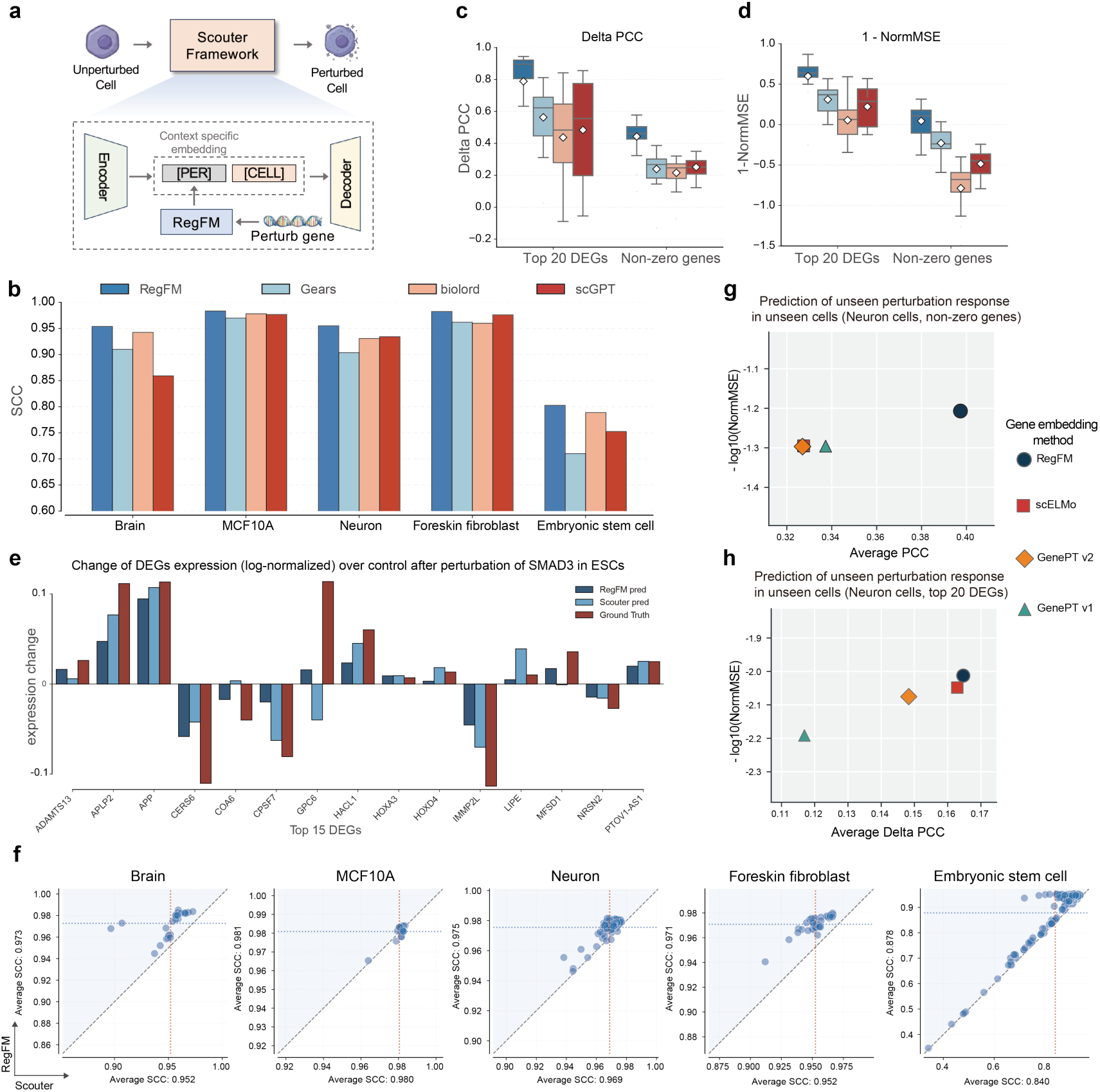
Fine-tuning RegFM to predict transcriptional response to genetic perturbation. **a,** Workflow for predicting the post-perturbation state from the corresponding unperturbed cell using the fine-tuned RegFM. **b,** SCC of gene expression between perturbed cells and control cells for RegFM and baseline methods across five datasets (Brain, MCF10A, Neuron, Foreskin fibroblast, and ESCs), evaluated on the top 20 DEGs. **c-d,** Delta PCC for predicted expression change (c) and 1 -NormMSE (d) for the top 20 differentially expressed genes (DEGs) and all non-zero expressed genes in post-perturbation prediction in embryonic stem cells (ESCs). **e,** Log-transformed predicted expression change of fine-tuned RegFM and observed expression changes for the top 15 DEGs in ESCs after genetic perturbation of *SMAD3*. **f,** Comparison of the fine-tuned RegFM and Scouter based on SCC for predicting post-perturbation expression changes across all non-zero expressed genes in the same five datasets. **g-h,** Comparison of fine-tuned RegFM with baseline methods for cross-dataset prediction of post-perturbation gene expression, evaluated on non-zero genes (g) and the top 20 differentially expressed genes (DEGs) (h). The *x* axis denotes PCC, and the *y* axis indicates −*log*_10_(*NormMSE*). The test perturbation gene set is derived from Neuron cells, which remains unseen during training.

Additionally, to evaluate the contribution of RegFM-derived gene embeddings in the Scouter framework, we compared them with GenePT^35^ derived gene embeddings under the same Scouter framework. As shown in **Fig. 4f**, across all five datasets, RegFM consistently improves performance in predicting all non-zero expressed genes, with higher average SCC observed in each dataset. For example, in the Foreskin fibroblast dataset, the average SCC increases from 0.952 to 0.971 when integrating GenePT embeddings with RegFM-derived embeddings (one-sided binomial test, *p*-value < 0.001). These results suggest that RegFM-derived omics-based gene embeddings are complementary to knowledge-based embeddings, can further improve predictive accuracy while preserving a lightweight model architecture.

Next, RegFM was further evaluated in a substantially more challenging setting that requires generalizing to both unseen gene perturbations and unseen cellular contexts. As shown in Supplementary Fig. S2c, RegFM generates context-specific gene embeddings from cell-type-specific trans-regulatory information, enabling cross-dataset prediction (see *Methods*). To this end, we performed a cross-dataset prediction experiment in which four datasets, excluding the Neuron dataset, were integrated as the training set. Three embeddings derived from Scouter were used as baseline methods for comparison^35, 42^. As shown in **Fig. 4h**, RegFM achieved a 6.0% improvement in PCC over the best baseline, GenePT v1, when predicting the transcriptional responses of the top 20 DEGs following 85 unseen perturbations in unseen neuron cells. A similar improvement was also observed when predicting the expression of non-zero genes (**Fig. 4g**). These results show that RegFM-derived embeddings provide useful information for modeling perturbational responses across perturbations and cellular contexts.

### RegFM reveals context-specific cis-trans regulatory interactions

In this section, we examined whether RegFM learns biologically meaningful relationships between cis-regulatory sequences, trans-acting regulators and target genes. As shown in **Fig. 5a**, the cross-attention module produces attention maps linking CREs to TF/CRs, thereby nominating context-specific regulatory relationships.

**Fig. 5.**
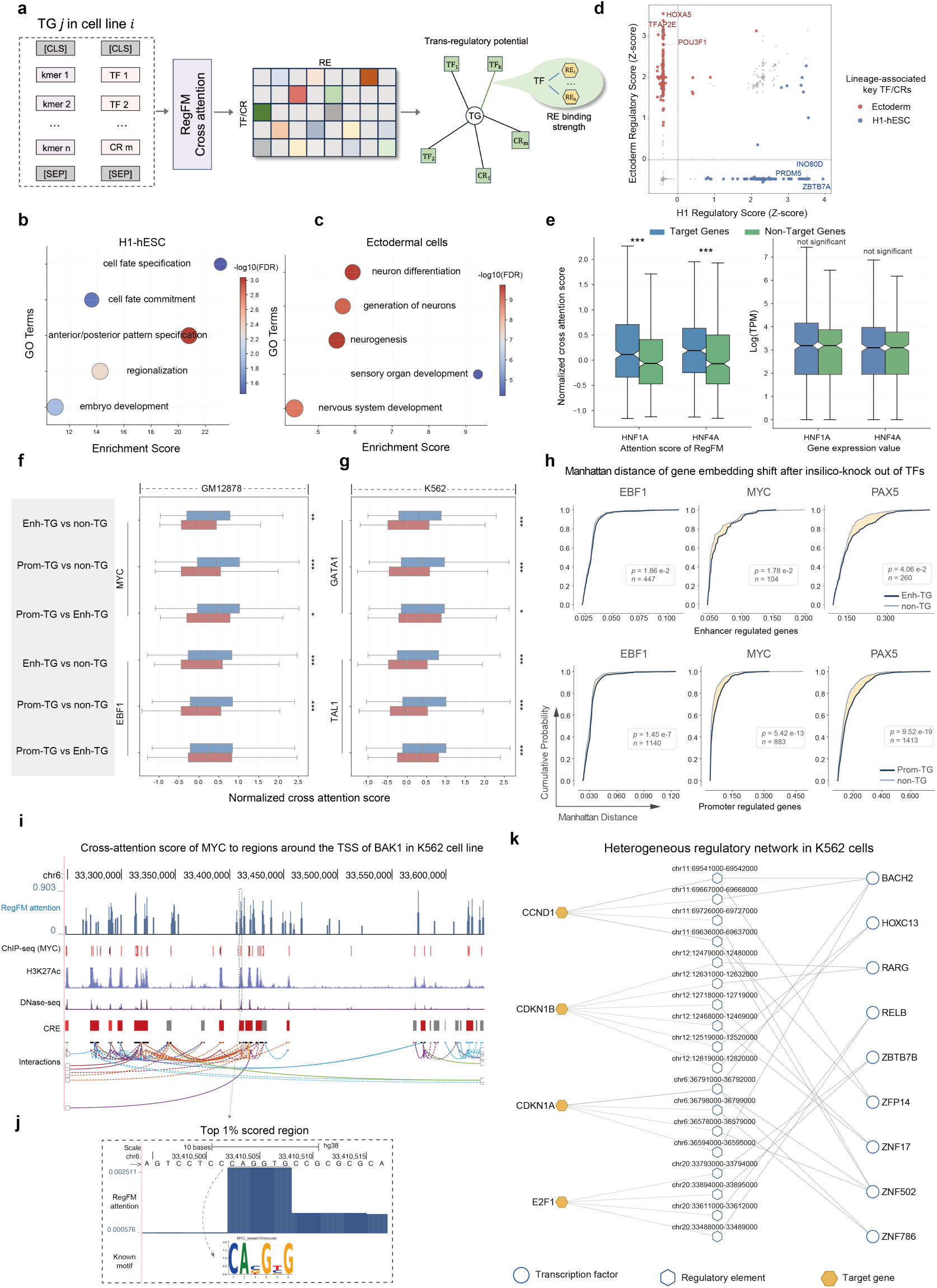
Interpretation and visualization of the cross-attention score between TF/CR and CREs. **a,** Workflow of RegFM for inferring regulatory strength between TFs and TGs, as well as between TFs and CREs. **b-c,** Gene ontology enrichment analysis of the top 100 attention-ranked TFs/CRs in H1-hESCs (b) and ectodermal cells (c), showing the top five cell-type-relevant terms in each context. **d,** Scatter plot of RegFM-derived regulatory scores in H1-hESCs (x-axis) and ectodermal cells (y-axis), where each point represents a TF or CR, blue indicates higher regulatory scores in H1-hESCs and red indicates higher regulatory scores in ectodermal cells. **e,** Box plots showing the distribution of attention scores for HNF1A, HNF4A on target genes and non-target genes defined by ChIP-seq in HepG2 (left), along with corresponding box plots of logarithm-transformed TPM values for the same gene sets (right). **f-g,** Box plots of attention scores for TFs in GM12878 (MYC and EBF1) and K562 (GATA1 and TAL1) across Enh-TGs, Prom-TGs, and non-TGs defined by integrated ChIP-seq and HiChIP-seq annotations, shown for GM12878 (f) and K562 (g), within each comparison, blue denotes Enh-TGs, Prom-TGs, and Prom-TGs, whereas red denotes non-TGs, non-TGs, and Enh-TGs, respectively. **h,** Cumulative distribution plots of Manhattan distances between gene embeddings under control and in silico knockout conditions for EBF1, MYC, and PAX5 in GM12878, with the x-axis representing Manhattan distance and the y-axis representing cumulative density; the upper panel compares Enh-TGs with non-TGs, and the lower panel compares Prom-TGs with non-TGs. **i,** Visualization of genomic tracks in the UCSC Genome Browser around the TSS of the BAK1 gene in K562, showing from top to bottom the RegFM-derived MYC attention scores across the region, MYC ChIP-seq, H3K27ac, DNase-seq, CRE annotations, and chromatin interactions between CREs. **j,** A representative example of the top 1% attention-scored regions in (i), where the highlighted region matches a known MYC binding motif. **k,** A showcase of the heterogeneous gene regulatory network in the K562 cell line inferred from the cross-attention matrix of RegFM. The network highlights the relationships among key genomic regions, TFs, and candidate target genes.

### Cross-attention scores reveal key regulatory TF/CRs

To assess the biological relevance of RegFM cross-attention maps, we analyzed human embryonic stem cell (H1-hESCs) and their differentiated ectodermal derivatives. For each cellular context, regions centered at TSSs of 200 HVGs were used as model input, and TF/CR importance was quantified by cross-attention scores averaged across layers, heads, and genomic regions (*see Methods*). Distinct sets of top-ranked TFs and CRs were observed between the two cellular contexts. Gene ontology analysis of the corresponding regulators showed enrichment for context-specific biological processes. In H1-hESCs, top-ranked TFs and CRs were associated with early developmental and cell fate-related programs (**Fig. 5b**), whereas in ectodermal cells, they were enriched for lineage-associated developmental processes, including organ and system development (**Fig. 5c**). Together, these findings suggest that RegFM-derived cross-attention scores reflect biologically meaningful, context-dependent regulatory signals. We next examined TFs and CRs showing differential regulatory activity between

H1 and ectodermal cells, focusing on those within the top 10% of ranked regulators in each context (see *Methods*). Regulatory scores were computed by z-score normalizing the aggregated cross attention scores. As shown in **Fig. 5d**, in ectodermal cells, the highest-ranked factors included established regulators of neuroectoderm and neural lineage specification, such as POU3F1^43^, members of the TFAP2 family^44^, and HOX family^45^, consistent with the inferred regulatory program. In H1 cells, top-ranked factors were enriched for regulators linked to pluripotency and chromatin-associated processes, including PRDM5^46^, ZBTB7A^47^. Collectively, these observations support that RegFM captures a coordinated shift from a pluripotent, chromatin-regulatory state to a lineage-specific ectodermal program.

### RegFM resolves TF-TG regulatory relationships

We next investigated whether cross-attention scores can reveal potential regulatory relationships between trans-regulators and target genes in a cell-type-specific manner. To quantify regulatory strength between TFs/CRs and target genes (TGs), we aggregated attention scores across all query positions and averaged across heads and layers, yielding a cumulative score that reflects the regulatory influence of query TFs/CRs on TGs. To assess the concordance between inferred regulatory strength and experimentally supported interactions, we leveraged ChIP-seq data. Specifically, we profiled HNF1A, and HNF4A in HepG2, key TFs involved in early cell fate specification and regulatory circuitry in human embryonic stem cells^48, 49^. Target and non-target genes were defined based on TSS distance-weighted ChIP-seq signal intensities (see *Methods*). As shown in Supplementary Fig. S3, RegFM assigns significantly higher attention scores to the top 500 protein-coding genes compared with the bottom 500 genes across all three TFs (one-sided Mann-Whitney U test, p < 0.01). Furthermore, integration with gene expression profiles showed that, even among genes without significant differences in expression, RegFM-derived attention scores still exhibited clear separation (**Fig. 5e**). These results suggest that cross-attention scores learned by RegFM capture TF-specific and cell-type-specific regulatory signals consistent with underlying biological regulation.

Next, we further validated RegFM at a more fine-grained level by integrating HiChIP-seq-derived chromatin interaction data with TF ChIP-seq signals. For each gene in each cell line, we retrieved corresponding chromatin interaction anchors from HiChIPdb^50^ and mapped them to the GRCh38 genome using liftOver^51^, and then integrated these anchors with TF ChIP-seq peaks to classify genes into three categories: enhancer-linked target genes (Enh-TGs), promoter-linked target genes (Prom-TGs), and non-target genes (non-TG). In GM12878 and K562 cell lines, we further evaluated the regulatory strength of MYC, EBF1, GATA1, and TAL1, all of which are known to play key regulatory roles in their respective cellular contexts^52–55^. As shown in **Fig. 5f-g**, predicted regulatory strength for both Prom-TGs and Enh-TGs was significantly higher than that for non-TGs across all TFs. Notably, for MYC, GATA1, and TAL1, RegFM-derived scores showed consistently higher regulatory strength for Prom-TGs compared with Enh-TGs, suggesting that cross-attention preferentially captures promoter-proximal regulatory interactions, which are typically more direct and constrained, whereas distal enhancer-mediated regulation is distributed across multiple regulatory elements and therefore exhibits weaker aggregate signal in the attention space.

### In silico perturbation of trans-regulatory sequence

Leveraging the trans-context transformer module, RegFM enables in silico knockout of specific TFs or CRs by modifying their activity states in the trans-regulatory input. We performed *in silico* knockouts of three key regulators EBF1, MYC, and PAX5 in the GM12878 cell line and quantified embedding shifts of candidate target genes under control and knockout conditions. For each gene, we computed the Manhattan (L1) distance between target gene embeddings under control and in silico knockout conditions as a measure of perturbation-induced representation shift. As shown in **Fig. 5h**, both Enh-TGs and Prom-TGs exhibited significantly larger embedding shifts than non-TGs as reflected by the rightward shifts of the empirical cumulative distribution curves. For example, among 1,413 candidate promoter TGs of PAX5, the average Manhattan distance reached 0.380, compared with 0.316 for non-TGs (one-sided Mann-Whitney U test, *p-value* = 1.05 × 10^−1^^6^). These results indicate that, in a zero-shot setting, in silico knockout of key TFs induces stronger perturbation effects in embedding space for their putative target genes, suggesting that RegFM captures context-specific TF-TG relationships and providing a foundation for systematic in silico perturbation analysis of gene regulatory programs.

### RegFM attention scores reveal context-specific CREs and regulatory networks

The cis-trans cross-attention module of RegFM allows us to derive TF-specific attention profiles across a genomic region. To assess whether these attention profiles reflect biologically meaningful TF-regulatory element interactions in a context-specific manner, we aggregated attention matrices across heads in the final cross-attention layer, yielding TF-specific attention profiles along genomic coordinates. In the K562 cell line, we focused on MYC, a master regulator of cell proliferation, and examined the genomic landscape surrounding the transcription start site (TSS) of the pro-apoptotic gene *BAK1*. As illustrated in **Fig. 5i**, across a distal span of more than 400 kb, the top 10% of high-attention regions identified by RegFM exhibited strong concordance with K562 epigenomic signals, including ChIP-seq binding peaks, H3K27ac histone modifications, DNase-seq accessibility signals, and annotated CREs. Furthermore, motif enrichment analysis via HOMER^56^ revealed that regions with higher attention were significantly enriched for high-scoring MYC motifs (Fig. S8b, one-sided Mann-Whitney U test, p = 0.0076). An example of MYC motif enriched in a top 1% attention region was visualized in **Fig. 5j**.

Furthermore, we constructed a context-specific heterogeneous gene regulatory network (GRN) based on the cis-trans cross-attention strength and the associated target gene (see *Methods*). As a demonstration example, we visualized the heterogenous GRN for the K562 cell line, integrating 9 TFs, top-associated genomic regions, and 4 candidate target genes (**Fig. 5k**). The inferred GRN showed biological interpretability. The predicted RARG-CDKN1B association was consistent with previous studies implicating RARγ signaling in p27-mediated cell-cycle arrest in leukemic cells^57^. Moreover, the network contained multiple ZNF-family transcription factors, a major class of regulators with established roles in hematopoietic regulation and cell-state-specific gene control^58–60^.

To interpret the features learned by RegFM, we derived sequence signatures (Position Weight Matrices, PWMs) from the cross-attention module and aligned them against the JASPAR database^61^ (see Methods section). Focusing on the K562 cell line, we examined the genomic regions of 13 genes associated with oncogenesis and proliferation, targeting 7 key TFs. RegFM successfully recovered TF-specific binding motifs with high fidelity. For instance, the similarity score for the GATA1 motif reached 0.437, significantly outperforming the background average of 0.186. A similarly robust recovery was observed for the MYB motif (Supplementary Figs. S8a).

In conclusion, these results demonstrate that RegFM prioritizes genomic regions with high TF-binding potential, confirming that its attention weights capture genuine biological syntax. By seamlessly integrating attention-based ranking and in silico perturbation, RegFM provides a highly interpretable framework for deciphering context-specific gene regulatory mechanism, highlighting the biological fidelity of the learned representations.

### Robustness analysis of RegFM

To assess the contributions of different modules in RegFM, we conducted extensive ablation experiments. First, we examined the contribution of masked language modeling pretraining for the two transformer modules. We therefore evaluated two ablated variants without pretraining in the gene expression prediction task. We found that self-supervised pretraining not only accelerated model convergence but also improved predictive performance, increasing the *R*^2^ from 0.447 without cis-DNA transformer pretraining to 0.492 (**Fig. 6a**). We next sought to investigate the contribution of the model architecture. To this end, we performed ablations on the two input modalities and conducted the same cross-cellular context gene expression prediction experiments. The results showed that the full RegFM model achieved the highest prediction performance in both the context-wise and gene-wise manner (one-sided paired Wilcoxon signed-rank tests *p*-value < 1e-3), highlighting the importance of integrating DNA sequence features and TF and chromatin regulator-associated gene expression signals for accurate modeling of gene expression (**Fig. 6b**). In the gene-wise setting, ablation of the regulation transformer abolished cross-cellular context predictive capacity, and the average *R*^2^ decreased from 0.232 in the full RegFM model to -0.043 (**Fig. 6c**). These results demonstrate that the regulation transformer module contributes not only to predictive accuracy but also plays a critical role in model generalization across cellular contexts. An ablation analysis was performed to evaluate the effect of sequence context length. Reducing the input length to 512 tokens led to decreases of 3.96% in average PCC and 5.88% in *R*^2^, indicating a potential benefit of longer sequence contexts for gene regulation prediction (Supplementary Fig. S10c).

**Fig. 6.**
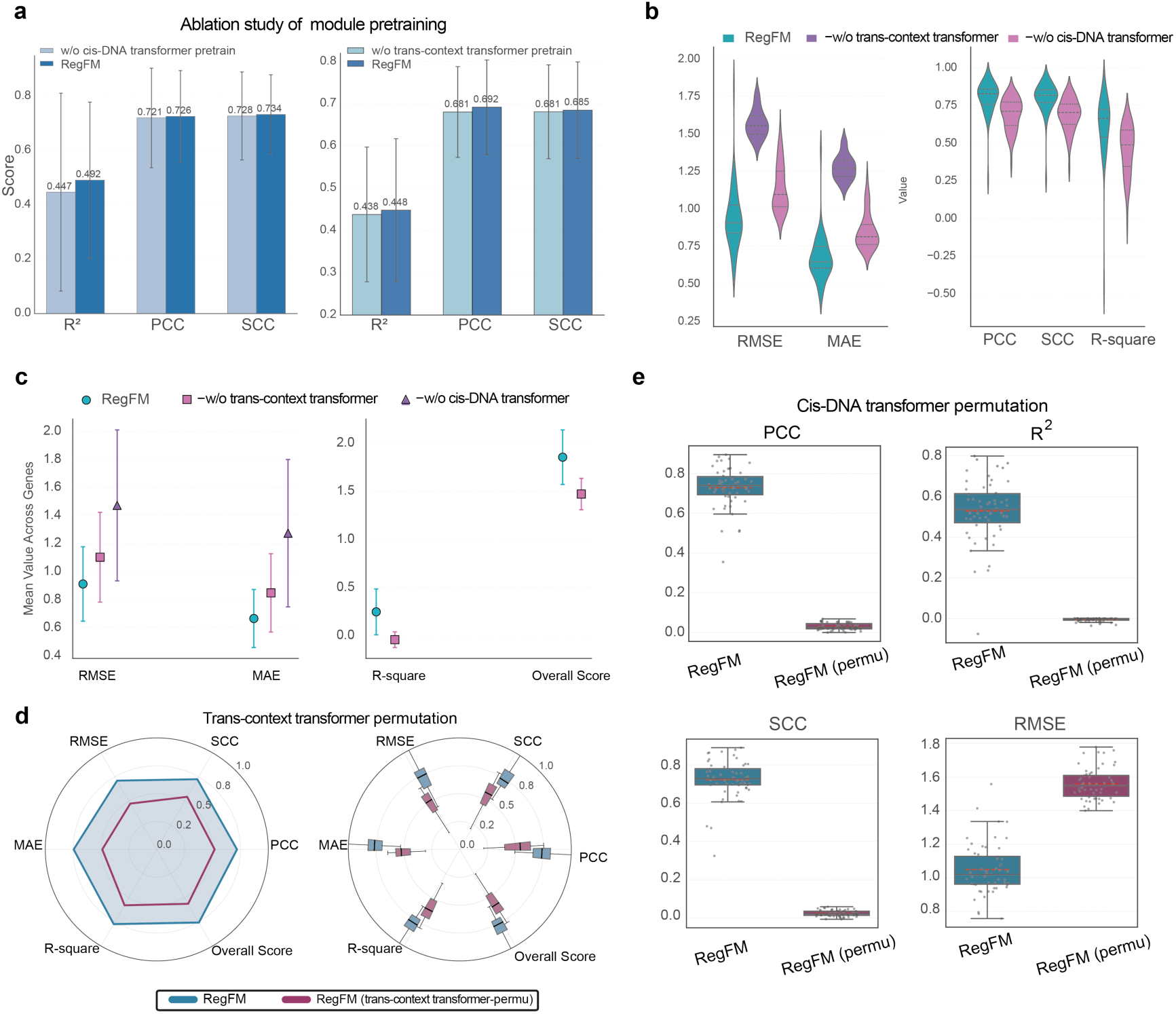
Ablation study of model architecture and training strategy of RegFM. **a,** The cross-cell-type gene expression prediction performance of RegFM and its variants without pretraining of the cis-DNA transformer and trans-context transformer modules. **b,** Comparison of gene expression prediction (context-wise) performance between RegFM and its architecture-ablated variants without the cis-DNA transformer and trans-context transformer modules. **c,** Comparison of gene expression prediction (gene-wise) performance between RegFM and its architecture-ablated variants without the cis-DNA transformer and trans-context transformer modules. **d,** Radar plot of cross-cellular-context gene expression prediction performance for RegFM and the trans-context transformer token permutation model (trans-context transformer-permu). **e,** Box plot of cross-cellular-context gene expression prediction performance for RegFM and the cis-DNA transformer token permutation model (cis-DNA transformer-permu).

We further examined whether the model correctly learned dependencies on token order. To this end, we introduced a random permutation strategy (see the *Methods*). As shown in **Fig. 6d-e**, permuting trans-regulatory tokens, as well as DNA k-mers, led to significant performance degradation (one-sided paired Wilcoxon signed-rank tests p-value < 1e-3). Notably, permutation of DNA k-mers resulted in a complete loss of the model’s ability to distinguish among different genes (**Fig. 6e**; Supplementary Fig. S10a-b). Collectively, these experiments validate the robustness of the core RegFM architectural design.

## Discussion

We present RegFM, a context-aware foundation model that provides a unified framework for modeling how genomic sequence is interpreted across cellular contexts. By coupling long-range cis-regulatory sequence with trans-regulatory state, RegFM moves beyond representations of DNA sequence or cellular state alone and enables transferable modeling of context-dependent transcriptional regulation. Across diverse predictive and interpretation analyses, our results show that the joint cis-trans representation supports broad context generalization while retaining biological interpretability. These findings establish context-aware cis-trans modeling as a promising direction for next-generation genomic foundation models.

Several directions could further extend the scope of RegFM. First, the current sequence modeling limited by variable-length k-mer tokenization. Extending RegFM to single-base resolution would enable more fine-grained regulatory modeling and better support applications such as regulatory variant interpretation. Second, the current RegFM primarily model focuses on gene expression prediction. Extending the RegFM to jointly model epigenomic features and other molecular signals would provide a more comprehensive representation of regulatory states and broaden its applications to multi-omics modeling. Third, incorporating naturally occurring genetic variation and experimental perturbation data could further extend RegFM toward causal regulatory modeling^62, 63^, enabling inference of context-specific regulatory effects and supporting more mechanistic and individualized interpretation of regulatory variation.

## Methods

### Data collection and processing

#### RNA-seq data collection and processing

To enable RegFM pretraining on gene regulatory patterns across a broad spectrum of cellular contexts, we systematically curated a large-scale collection of bulk RNA-seq data from the ENCODE consortium^6^ and single-cell RNA-seq datasets from CellxGene^7^. For the bulk RNA-seq data, we collected 1,018 RNA-seq experiments aligned to the GRCh38 reference genome from the ENCODE consortium (Supplementary Table S1 and S3, Fig. 1a). We extracted TPM values for protein-coding genes and firstly applied a log transformation. For the expression level *e*_*i*_ of gene *i* (TPM value), the transformation can be formulated as 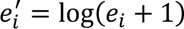. We then aggregated the log-transformed TPM values according to the ‘Biosample term’, averaging replicates derived from the same cell line or tissue, resulting in a total of 281 distinct cellular contexts. Gene expression profiles across these contexts were then subjected to quantile normalization, resulting in a cellular-context-by-gene matrix X_B_ ∈ *R*^*c* × *m*^, where *c* denotes the number of cellular contexts and *m* denotes the number of protein-coding genes. For the single-cell RNA-seq data, we collected approximately 43 million cells from CellxGene, selected 30 million normal cells, and then aggregated single-cell gene expression read counts into pseudo-bulk data across 663 cell types. Similar with the procedure used in GET^5^, we then applied a logarithmic transformation to the read counts, resulting in a cell type-by-gene matrix X_S_ ∈ *R*^*k*^ ^×^ ^*n*^, where *k* denotes the number of cellular contexts and *n* denotes the number of protein-coding genes. A comprehensive summary of the pseudo-bulk cell types employed for gene expression prediction is presented in Supplementary Table S2.

#### Transcription factors and chromatin regulators

To comprehensively model the regulatory landscape across diverse cellular contexts in RegFM, we curated datasets of transcription factors (TFs) and chromatin regulators (CRs). Specifically, we collected 1,639 TFs from Cis-BP^25^ and 563 CRs from CRdb^26^, and after deduplicating the list, we obtained a final set of 2,103 unique TFs and CRs, which constitute the input “sentences” for the regulatory transformer pretraining of RegFM. A detailed table containing information on these TFs and CRs is available in Supplementary Table S4.

### Model architecture of RegFM

The overall architecture of RegFM consists of three core modules: a cis-regulatory sequence encoder (cis-DNA transformer), a trans-regulatory context encoder (trans-context transformer), and a cross-attention module that couples the two pathways. The cis-DNA transformer module processes DNA sequences within a broad gene-centered regulatory window, which are tokenized into variable-length, non-overlapping k-mers via byte-pair encoding, and learns the inherent regulatory grammar embedded in cis-regulatory elements. The trans-context transformer module encodes dynamic cellular contexts by modeling the activity states of TFs and CRs, capturing their context-dependent expression and functional profiles. These two complementary representations are then integrated through the cross-attention mechanism, which explicitly models context-specific interactions between trans-acting regulators and cis-regulatory sequences. By dynamically weighting relevant sequence patterns according to the availability and activity of cognate regulators, the model enables context-aware interpretation of gene regulation and ultimately determines the resulting gene expression levels.

#### Cis-regulatory sequence tokenization

To reduce sequence length and improve sample efficiency, we adopt SentencePiece^64^ with Byte Pair Encoding (BPE) for DNA sequence tokenization^13^. Unlike overlapped k-mer tokenization or using base pairs as tokens, BPE tokenization segments DNA sequences into non-overlapping units to represent a given DNA region. Beyond compressing sequence length, distinct k-mers as minimal units may capture TF binding motifs, thereby enabling the model to more effectively learn regulatory relationships between TFs and CREs. From a computational efficiency perspective, a larger vocab size enables more efficient compression of information in long-range DNA regions. Therefore, we split the whole GRCh38 reference genome to non-overlapping regions and trained a BPE tokenizer, with the vocab size of 150, 000, which lies between 4^8^ and 4^9^ and allows coverage of the majority of k-mers with lengths up to 10. During pretraining for gene expression prediction, tokenized input sequences are truncated to the 71,680 whenever the tokenized sentence exceeds this limit. A detailed analysis of the maximum input sequence length in RegFM is provided in Supplementary Fig. S3.

#### Cis-regulatory transformer module

To enable RegFM to perceive a broader DNA sequence context and thereby capture potential distal regulatory interactions, we trained a DNA transformer module with long-range DNA sequence inputs, using high-resolution k-mer tokens for sequence representation. As the core of RegFM, the DNA transformer module comprises 6 LongNet^65^ encoder layers with a bidirectional multi-head dilated attention mechanism to model global dependencies in long-range DNA regions surrounding the TSS of genes, thereby capturing regulatory networks across diverse cellular contexts. Following the DNA tokenization, the ± 325 kbp DNA regions centered at the TSS of gene *j* are segmented into a kmer sentence, denoted as *C_j_* = (*K*_1_, *K*_2_, …, *K_n_j__*). Two special tokens, [CLS] and [SEP], are appended to the start and end of the k-mer sequence, thereby yielding 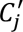. Then, the embedding layer takes the k-mer sequence as input and outputs a word embedding for each k-mer token. In addition, a trainable weight matrix *W*_*p*_ maps the absolute position of each kmer token to a positional embedding, which is then added to the word embedding to form the input embedding for the LongNet encoder. The output of the embedding layer can be formulated as

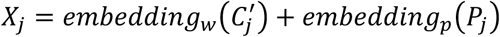

where embedding_w_ denotes the word embedding layer, embedding_p_ denotes the positional embedding layer, and *P_j_* denotes the absolute positional encoding. The input embedding dimension for each token is set to 768.

Each transformer block operates on inputs and outputs with an embedding dimension of 768 and uses 12 attention heads. Based on dilated attention mechanism, the input and output of each single LongNet encoder block can be formulated as:

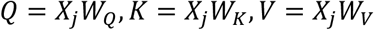

where *W_Q_*, *W*_*k*_, and *W*_*V*_ are trainable weight matrices used to compute the Query (Q), Key (K) and Value (V), respectively. To improve computational efficiency, dilated attention is incorporated into the self-attention mechanism^66^. Specifically, (*Q*, *K*, *V*) are partitioned into segments of length *w*, and within each segment, attention is computed in a sparsified manner with an interval *r*. The computation within each segment can be formulated as follows^65^:

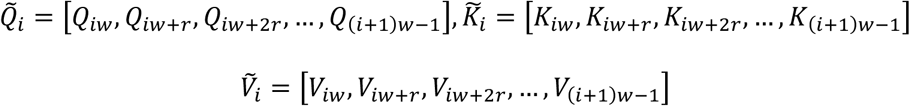

The computations of these sparse segments are performed in parallel, and the results are concatenated to produce the final output:

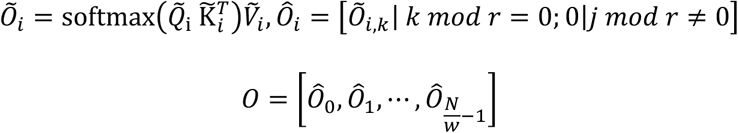

where *N* denotes the sequence length. The outputs of all heads are then concatenated to form the final output. The proposed multi-head dilated attention mechanism substantially improves computational efficiency in terms of both runtime and memory for long sequences. During training, we used distinct segment lengths and corresponding dilation rates, specifically [(128, 16), (512, 64), (1024, 256), (2048, 512)].

#### Trans-regulatory transformer module

The regulation transformer module is designed to endow RegFM with cellular context awareness, enabling it to learn context-specific expression patterns of TFs and CRs across diverse cellular contexts. Following the RNA-seq data processing pipeline, we first discretized the normalized TPM values of all protein-coding genes across 1,018 bulk RNA-seq experiments into 256 bins based on the global distribution. This yields a discretized expression value for each gene in each cellular context. As the input sentence to the regulation transformer module, *E*_*i*_ = (*Ex*_1,*i*_, *Ex*_2,*i*_, …, *Ex*_*m*,*i*_) denotes the discretized expression values of *m* selected TFs and CRs in the *i*^th^ cellular context, where each cellular context corresponds to a cellular sentence, and *m* is set to 2103. Two special tokens, [CLS] and [SEP], are also appended to the start and end of the cellular sequence, thereby yielding 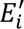.

Taking *E_i_* as the input sentence, the embedding module of the regulation transformer consists of two components: an expression embedding layer and a regulator embedding layer. Discretized regulator expression values are treated as tokens and mapped to expression embeddings through the expression embedding layer. The regulator embedding layer takes the ordered sequence of regulators *R* = (*R*_1_, *R*_2_, …, *R*_*m*_) as input and produces regulator-specific embeddings. As the ordering of CRs and TFs within the input sequence is fixed across cellular contexts, the regulator embeddings inherently encode positional information and thus serve the role of positional embeddings. The final input representation to the regulation transformer is obtained by summing the expression embeddings and the corresponding regulator embeddings. The regulation transformer is implemented as a stack of transformer encoder blocks, consisting of 12 layers with 6 attention heads per layer.

#### Cross-attention module

To enable RegFM to learn cell-type-specific regulatory mechanisms between regulatory elements proximal to the TSS and genes from large-scale data, rather than being restricted to knowledge curated in existing databases, we align the output sequences *O_DNA_j__* from the DNA transformer and the output sequences *O_Reg_i__* from the regulation transformer via cross-attention mechanism (**Fig. 1**). Specifically, taking *O_DNA_j__* as the query and *O_Reg_i__* as the key and value, the cross-attention computation at each stacked layer can be formulated as follows:

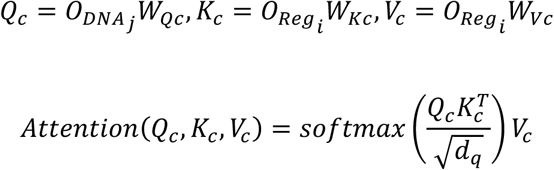

where *d*_*q*_ denotes the dimensionality of the query latent embedding, *j* denotes the *j*^*th*^ gene, and *i* denotes the *i*^*th*^ cellular context. The computation of multi-head attention follows the same formulation as described in the DNA transformer module, and the outputs of all heads are concatenated to form the final output. In RegFM, we employ 4 cross-attention layers, each with two attention heads. To predict the normalized expression value of a gene, we extract the [CLS] token from the output sequence of the cross-attention module as a sentence-level embedding and use it as the input to a regression head. This prediction head consists of a fully connected layers, with a *LeakyReLU* activation applied at the output, to perform regression-based prediction of gene expression values. The process can be formulated as:

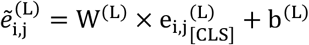

where *W*^(*L*)^ and *b*^(*L*)^ are learnable parameters of the *L*^*th*^ layer in prediction head, and 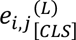 denotes the cellular-context specific gene embedding. In summary, the complete model comprises 302 million trainable parameters.

### Training strategy for RegFM

To enable RegFM to comprehensively capture the biological context of the human genome and transcriptome, as well as the cis-regulatory relationships between CREs and genes and the trans-regulatory relationships between TFs/CRs and genes, we apply three pretraining tasks. These include masked language modeling for the DNA sequence module, masked language modeling for the regulator sequence module, and a cross-cellular-context gene expression prediction task. Pretraining is conducted in a staged manner, where the three tasks are applied sequentially across successive training stages.

**(i) Self-supervised pretrain stage.** We will pretrain both the cis-DNA transformer (cis-T) and trans-context transformer (trans-T) via self-supervised strategy using the massive amount of unlabeled genomic data. **Masked language modeling of cis-DNA transformer module.** Using the human GRCh38 reference genome, we first excluded blacklisted regions^67^ on the reference genome and partitioned the remaining genome into 200kbp regions. These regions were further generated using a sliding window with a stride of 50 kbp. On these regions, we applied a masked language modeling (MLM) pretraining strategy at the sentence level, in which approximately 5% of the tokens in each input sentence were randomly masked and predicted using an additional prediction head. The model was trained using a distributed data parallel (DDP) strategy on 8 NVIDIA RTX A6000 GPUs, each with 48 GB of memory. The batch size was set to 64, with a gradient accumulation step of 12. **Masked language modeling of trans-context transformer module.** Analogous to the DNA transformer module, we performed pretraining on regulator sequences, where the TF/CR expression patterns within each cellular context were represented as a sentence. A MLM pre-training task with a masking ratio of 15% was applied across 1,018 experiments. Details of the ablation studies assessing the impact of trans-context transformer module pretraining on gene expression prediction performance are provided in Fig. 6.
**(ii) Supervised train stage.** After pretraining, we performed supervised training to map paired inputs (cis-sequence and trans-context) to measured molecular phenotypes (e.g., gene expression) using public resources such as ENCODE. Note that each training instance corresponds to a genomic region and a cellular context pair. This design yields substantially more effective training instances than sequence-only models since a single locus is observed under various regulatory contexts. **Cross cellular context gene expression prediction task.** To enable the cross-attention layers to effectively learn cell-type-specific cis-and trans-regulatory relationships, we performed pretraining using protein-coding genes from 281 cell lines and tissues curated from ENCODE, together with 663 cellular contexts from CellxGene. Given a 650 kbp region centered at the TSS of a gene and the corresponding regulator sentence in a specific cellular context as inputs to RegFM, the pretraining objective is to regressively predict the gene’s expression level in that cellular context. The mean squared error (MSE) loss is used for optimization. RegFM is implemented using the PyTorch framework^68^ and trained with a DDP strategy^69^ on 8 NVIDIA RTX A6000 GPUs. Training is performed with a batch size of 64, using the Adam optimization algorithm^70^ with a learning rate of 1e-5 and a gradient accumulation step of 8.
**(iii) Fine-tune stage.** After the first two stages of training, RegFM is able to capture cellular context-specific representations of genomic regions. Building upon these embeddings, a wide range of genome functional annotation tasks can be accomplished by feeding the sequence embeddings into newly introduced prediction heads, including cell-type-specific regulatory element prediction, gene dosage sensitivity prediction, promoter bivalency prediction, genetic perturbation response prediction, and the inference of gene regulatory relationships through the interpretation of cross-attention weights.

### Downstream analysis

#### Gene expression prediction in unseen cellular contexts

To comprehensively evaluate the performance of RegFM in modeling gene expression across diverse cell types, we conducted cross-cell-type gene expression prediction. All predictions were derived from sentence-level genomic region embeddings (the [CLS] token) produced by the cross-attention module. Let *Y*^pred^ ∈ ℝ^m×n^ and *Y*^true^ ∈ ℝ^m×n^ denote the predicted and ground-truth gene expression matrices, respectively, where m represents the number of unseen cellular contexts in the test set and n denotes the number of selected genes (for example, highly variable genes or all protein-coding genes). Two complementary evaluation strategies were employed. (a) Context-wise manner: for each cellular context *i* ∈ {1, …, *m*}, performance was quantified by computing evaluation metrics based on the similarity between the predicted and true gene expression vectors 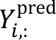 *and* 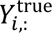, thereby assessing the model’s ability to capture gene-wise expression variation within unseen cell types. (b) Gene-wise manner: for each gene *j* ∈ {1, …, *n*}, metrics were computed based on the similarity between the predicted and true expression vectors across cellular contexts, 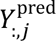 *and* 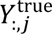, enabling evaluation of the model’s ability to capture cell type-specific expression patterns for individual genes. Together, these two evaluation schemes provide a comprehensive assessment of the model’s capacity to characterize differential gene expression across both cellular contexts and genes.

For the gene expression prediction task, the fine-tuned RegFM was compared with eight alternative methods: DNABERT, Nucleic Transformer, Nucleotide Transformer, DNABERT-2, EVO-2, Xpresso, LINGER, and GET. All methods were implemented following their respective published studies and publicly available GitHub repositories. For models with available pretrained weights, we initialized from the released checkpoints and fine-tuned them on the same training data as RegFM. Notably, DNABERT provides four pretrained variants (4-mer, 5-mer, 6-mer, and 7-mer), we report results based on the 6-mer model, as no significant performance differences were observed among variants for this task (Supplementary Figs. S4). For EVO-2, the 7B backbone parameters were frozen, and only a newly introduced prediction head was fine-tuned. In the inference implementation released in the EVO-2 GitHub repository, a base-level embedding *e*_*i*_ is generated for each nucleotide position. We extracted 8,192 tokens centered around the TSS, denoted as 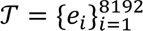, and summed all embeddings within this region to obtain a region-level representation, 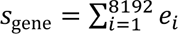. This gene embedding was subsequently fed into a prediction head for downstream task-specific prediction. For the gene expression task-specific model Xpresso, we used the version based solely on DNA sequence and trained it from scratch on the same data. Third, for methods requiring paired data, a subset of the dataset was selected for training and comparison. Specifically, LINGER requires chromatin accessibility data as paired input and is a neural network-based approach that trains a separate model for each gene. Since LINGER is not a pretrained foundation model, both LINGER and RegFM were trained from scratch using bulk data to ensure a fair comparison. DNase-seq and ATAC-seq data aligned to the GRCh38 reference genome were obtained from ENCODE. Notably, we relaxed the pairing criteria at the cell line or tissue level, considering RNA-seq and chromatin accessibility data as paired when both were available within the same cellular context. Using this approach, we obtained 26 cellular contexts with paired ATAC-seq data and 111 cellular contexts with paired DNase-seq data. For the GET model, the published pretrained model was used directly for comparison with RegFM. RegFM was trained from scratch on the PBMC dataset, with each leave-one-out cell type selected as the test cellular context, following the same settings as those described in the original GET model release tutorial on GitHub. It is important to note that the training data for RegFM consisted solely of a subset of the data used to pretrain GET, thereby creating a comparison setting that is more favorable to GET model.

#### Cis-regulatory element classification

In the CRE identification task, RegFM was fine-tuned by initializing the trans-context transformer module with the pretrained weights and feeding the [CLS] token representation from the cross-attention module into a newly introduced classifier for both binary and multi-class classification. All model parameters were jointly fine-tuned without freezing any weights during training. The classification layer is a two-layer fully connected neural network with 256 units in the hidden layer. Additionally, in the CRE classification task, the input to the cis-DNA transformer module is fixed at 300 bp for binary classification and 1200bp for multiple classsification, as most typical cis-regulatory elements are defined as short DNA sequences, typically ranging from 100 to 1,000 bp^71^. With this shortened DNA sequence length, the vocabulary for tokenization in the cis-DNA transformer module of fine-tuned RegFM is adapted to the k-mer vocabulary of DNABERT-2, which includes 4,096 k-mers, enabling better adaptation to the tokenization of short DNA sequences. For the comparison, a total of eight baseline methods were applied to this task: DNABERT, DNABERT-2, DeepSEA, RF, SVM, NT, Logistic Regression (LR), and CREATE. DeepSEA^72^ is a proprietary neural network method used for predicting the functional and epigenetic properties of genomic regions. CREATE^4^ is a CRE identification method based on VQ-VAE^73^ and multi-omics data, and was specifically included for comparison in the insulator identification task using samples with matched multi-omics data. For the 3 traditional machine learning classifiers, we used the k-mer frequency vector as the input feature for the classifier. This feature is commonly used to extract encoding information from DNA sequences and serves as an input feature for computational methods^74, 75^. The k-mer frequency vector is computed as:

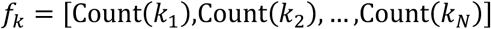

where *f*_*k*_ is the k-mer frequency vector, *k*_1_, *k*_2_, …, *k*_*N*_ represent all possible k-mers in the given DNA sequence, and Count(*k*_*i*_) is the frequency of each k-mer *k*_*i*_ within the sequence.

The annotation data for CREs were sourced from two datasets. The first is the CRE annotation from EnhancerAtlas 2.0^28^, which provides annotated CRE regions across 54 cell lines/tissues. After pairing with RNA-seq data based on cell types, we obtained enhancer annotations for 44 cell lines/tissues. Notably, the background regions were derived from DNase-seq peak files from ENCODE (Supplementary Table S6) corresponding to the relevant cell lines/tissues. The enhancer regions within the respective cell lines were excluded, and the remaining regions were considered as background regions, thus forming the negative samples. The second dataset, sourced from Cui et al.^4^, provides multi-class element annotations based on ChIP-seq and DNase-seq data, covering the cell lines HepG2, GM12878, K562, and Hela-S3, as well as three types of regulatory elements (Enhancer, Insulator, Silencer) and background regions. Since previous promoter prediction methods can already achieve accuracies exceeding 0.9, we excluded promoters from the multi-class categorization. The silencer elements in this dataset are sourced from SilencerDB^76^.

The primary focus of this task is the cell-type specificity of various regulatory elements, aiming to capture distinct regulatory patterns across different cell types. To comprehensively evaluate the model’s ability to distinguish cell-type-specific CREs, we employed three different data partitioning strategies: Setting (i): We combined CRE annotations from multiple cell lines and randomly partitioned the data into training and testing sets, without distinguishing cell-type-specific samples. Setting (ii): We integrated CREs from all cell lines into a common library and selected cell-type-specific CREs (i.e., a CRE region may have different functions across cell types and thus belong to different CRE categories). This resulted in a CRE-by-cell line matrix *C*^*N_cre,_N_cell_*^, where *N*_*cre*_ represents the number of CRE types and *N*_*cell*_ the number of cell types. The pairs of CRE labels and cell lines were mixed, and the matrix was flattened, followed by random partitioning into training and testing sets to assess the model’s ability to differentiate cell-type-specific CREs. Setting (iii): We partitioned the data by cell type into training and testing sets to evaluate the model’s generalization ability in identifying CREs in unseen cell lines. To evaluate classification performance, we primarily used the auROC, auPR, F1 score, and accuracy as metrics. For the multi-class task, the macro-averaged method was employed to assess the model’s performance across all categories.

#### Promoter bivalency prediction

In the promoter bivalency prediction task, the fine-tuned RegFM adopted a fine-tuning strategy similar to that used in the CRE classification task. The key distinction is that only the newly introduced classification head was trainable, while the backbone parameters of RegFM remained frozen throughout the fine-tuning stage. The annotation data for bivalently marked genes, as well as genes that are unmethylated or marked solely by H3K4me3 or H3K27me3, were sourced from a previously reported study^30^. This gene set comprises 332 genes in embryonic stem cells, with promoter states annotated for each gene. Bivalently marked genes were labeled as positive samples, whereas all remaining genes were treated as negative samples, thereby enabling evaluation of the ability of RegFM and baseline methods to distinguish bivalently marked genes. In this experiment, RegFM was fine-tuned using TF/CR gene expression reference data from embryonic stem cells obtained from ENCODE as the cellular context.

For a fair comparison of the classification performance achieved using gene embeddings derived from fine-tuned RegFM and baseline methods, we included two representative baseline approaches capable of generating genomic region embeddings: GenePT^35^ and EVO-2^16^. These methods represent two complementary sources of prior information, namely knowledge-driven embeddings derived from textual annotations (e.g., gene descriptions from official National Center for Biotechnology Information) and sequence-derived embeddings learned from large-scale omics data, respectively. Using the pre-trained RegFM, GenePT, and EVO-2 models, an embedding vector was obtained for each gene. A key distinction is that the embeddings produced by RegFM are specific to embryonic stem cells (ESCs), as they are conditioned on the corresponding cellular context, whereas the baseline methods generate general-purpose gene embeddings that are shared across different cellular contexts. For classification, each gene embedding was fed into a multilayer perceptron (MLP) consisting of two linear layers, followed by a sigmoid activation function to perform binary classification. The gene set was evaluated using 5-fold cross-validation.

#### Gene dosage sensitivity prediction

In the gene dosage sensitivity prediction task, the fine-tuned RegFM adopted the same fine-tuning strategy as used in the previous task, with the RegFM backbone weights remaining fixed throughout training. A key difference is that the gene set in this task is not associated with a specific cellular context. Therefore, to comprehensively characterize the regulatory function of each gene across diverse cellular contexts, 281 cell lines and tissues from ENCODE were used as reference cellular contexts, and an attention-based fusion module was applied to integrate gene embeddings derived from these distinct contexts before being fed into the classification head. For each gene *i*, RegFM generated a set of context-specific embeddings across the *N* reference cellular contexts:

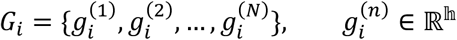

where ℎ denotes the hidden dimension. For a gene set containing *m* genes, this yields a representation tensor *G* ∈ ℝ^m×ℕ×h^. To integrate regulatory signals across cellular contexts, we applied an attention-based fusion module to model pairwise dependencies among context-specific embeddings:

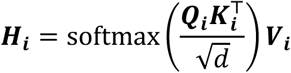

where *Q*_*i*_, *K*_*i*_, *and* *V*_*i*_ denote the query, key, and value projections of *G*_*i*_, respectively. The final fused representation was then obtained by pooling across the context dimension: *g̃_*l*_* = Pool(*H_i_*) ∈ ℝ^*h*&^, thereby capturing the global regulatory characteristics of each gene across diverse cell types in a more comprehensive manner than any single-context representation alone. The fused embedding was subsequently concatenated with the sequence-derived representation and fed into a two-layer fully connected classification head for downstream prediction.

Gene labels were curated from a previously published benchmark study on dosage-sensitive genes^31^. Following the original definition, genes classified as the most reliable dosage-sensitive (MRDS) were treated as positive samples, whereas genes classified as the most reliable dosage-insensitive (MRDIS) were used as negative samples. Briefly, MRDS genes were defined as those consistently identified across four independently curated dosage-sensitive gene datasets, while MRDIS genes were defined as genes absent from all four datasets and with loss-of-function intolerance probability (pLI < 0.5) in both haploinsufficiency-based resources. This dataset was subsequently used to formulate a binary classification task for evaluating the ability of RegFM and baseline methods to distinguish dosage-sensitive genes.

#### Genetic perturbation transcriptional response prediction

For the genetic perturbation response prediction task, RegFM was used to generate reference gene embeddings that capture context-aware regulatory representations. These embeddings were subsequently incorporated into the variational autoencoder (VAE) framework in Scouter to improve the prediction of gene expression responses genetic perturbations. **Fig. 4a** and **Supplementary Fig. S2** illustrates the model architecture used for this downstream task. The expression profile of a control cell is first passed through a compressor to obtain a 64-dimensional cell-state vector. Meanwhile, the cis-regulatory sentence derived from the ±325 kbp DNA region surrounding the perturbed gene and the corresponding cell-type-specific trans-regulatory sentence are fed into the cis-DNA transformer and trans-context transformer modules of RegFM, respectively. Through the cross-attention module, these inputs are integrated to generate a 768-dimensional perturbation embedding. The concatenation of the cell-state vector and the perturbation embedding is subsequently fed into a generator to predict the corresponding transcriptional response. In addition, we further developed an attention-fusion variant that integrates the RegFM-derived perturbation embedding with the perturbation embedding from Scouter, thereby further improving predictive accuracy (Supplementary Text S3).

RegFM is fine-tuned using the same objective function as Scouter, which consists of an autofocus loss and a direction-aware loss. For a batch of *N* triplets 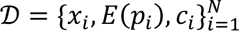, where *x*_*i*_ denotes the cell embedding of perturbed cell *i*, *p*_*i*_ denotes the perturbed gene, *E*(*p*_*i*_) denotes the perturbation embedding generated by RegFM, and *c*_*i*_ denotes the cell embedding of the corresponding control cell, the generator predicts the post-perturbation embedding *x̂*_*l*_. The autofocus loss is defined as:

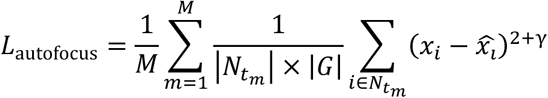

where|*N_t_m__*| denotes the number of triplets associated with perturbation *t*_*m*_, |*G*| denotes the dimensionality of the embedding vector, and *γ* controls the extent to which larger prediction errors are emphasized. This loss encourages accurate reconstruction of the perturbed cellular state while placing greater weight on large discrepancies. In addition, the direction-aware loss defined as:

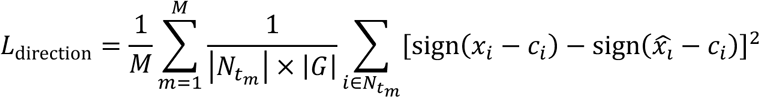

which explicitly penalizes inconsistencies between the predicted and observed directions of change with respect to the control cell embedding. The final optimization objective is formulated as *L* = *L*_autofocus_ + *λL*_direction_, where *λ* balances reconstruction fidelity and directional consistency. To predict the response to a perturbation on gene g, we adopted a strategy similar to that used in Scouter, in which *K* randomly selected control cells were used as inputs, and the estimated response was obtained as 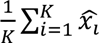, a strategy that has been reported by Scouter^32^ to yield robust predictions, and *K* is set to 300 here.

For comprehensive comparison, four baseline methods were included: GEARS^34^, biolord^33^, scGPT^21^, and Scouter^32^. Notably, Scouter provides three distinct perturbation gene embedding libraries, namely GenePT v1, GenePT v2, and scELMo^42^, which are derived using different versions of LLMs and textual prompts to encode perturbation genes. All baseline methods were implemented following their original architectures and input formats, strictly adhering to the tutorials and protocols provided in their respective GitHub repositories. For scGPT, we likewise used its released pretrained model and followed the official GitHub tutorial for perturbation response prediction. The discrepancy between our results and those reported in their study is likely attributable to differences in software versions or subsequent updates.

A total of five processed datasets from the PerturBase database were used for evaluation, corresponding to five distinct cell types: Brain^37^, MCF10A^38^, Neurons^39^, embryonic stem cells^40^, and foreskin fibroblasts^41^. For within-dataset prediction, 20% of each dataset was held out as the test set, while the remaining 80% was further split into 90% for training and 10% for validation. The validation set was used for early stopping, and, in addition, all perturbation genes within each dataset were partitioned using a 5-fold cross-validation scheme to provide a more comprehensive benchmark of the predictive performance of the fine-tuned RegFM and baseline methods.

To evaluate model performance, we employed four metrics, focusing on the top 20 DEGs and non-zero expressed genes, as not all genes exhibit substantial expression changes before and after perturbation. Given the expression vector of a perturbed cell *x*_*i*_, a control cell *c*_*i*_, and the prediction *x̂*_*l*_, the metrics are defined as:

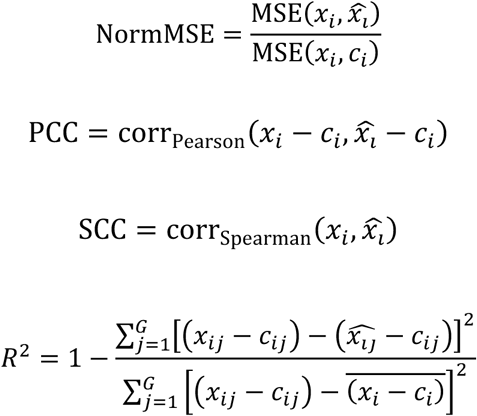

where *d_j_* denotes the rank difference between *x*_*i*j_ and 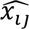 and *G* denotes the number of genes under evaluation. Collectively, these metrics assess not only the accuracy of predicting post-perturbation expression levels, but also the ability to capture the directionality of perturbation-induced expression changes.

Furthermore, for cross-dataset or -condition prediction, we evaluated a leave-one-out strategy in which datasets were integrated as follows: (1) using Scanpy^77^, genes were aligned across datasets by taking the intersection of shared genes as anchors; (2) batch effects were corrected using Harmony^78^; (3) for data splitting, the neuron cell dataset was designated as the test set, while the integrated dataset composed of the remaining four datasets was used for training. All remaining procedures were kept consistent with the single-dataset prediction setting. For GenePT and scELMo, the gene embeddings are dataset-independent and therefore do not require explicit dataset labels. In contrast, for RegFM, overlapped genes across different datasets were annotated with their corresponding cell-type identities, and the reference gene embeddings derived from ENCODE were used to represent each gene. As illustrated in **Fig. 4b**, the overall model framework remains consistent with the single-dataset setting, with the key distinction that the fine-tuned RegFM explicitly distinguishes the cell-type context of perturbed genes across different datasets.

#### Cross-attention score analysis

Given that RegFM takes the cis-regulatory sequence as the query and the trans-regulatory sequence as the key and value in cross-attention module, it enables characterization of the regulatory strength of trans-acting regulators on target genes, the attention to regulatory elements, and the context-specific importance at the cellular level.

#### Context-specific regulatory scores of trans-acting regulators

To derive regulatory scores of trans-acting regulators in a specific cellular context, we first selected the top 200 HVGs based on processed gene expression values in that context relative to others. For each gene, the cis-regulatory sequence *C_j_* = (*K*_1_, *K*_2_, …, *K_n_j__*) and the trans-regulatory sequence *R*_*i*_ = (*R*_1_, *R*_2_, …, *R_m_i__*), corresponding to the *j*-th gene in the *i*-th cellular context, were used as inputs to RegFM. After averaging the attention matrices from the cross-attention module across layers and heads, we obtained an attention *A*_*i*j_ ∈ ℝ^ℕ_s_×L_c_×L_r_^, where *N*_*s*_ denotes the number of selected HVGs, *L*_*c*_ the length of the cis-regulatory sequence, and *L*_*r*_ the length of the trans-regulatory sequence. We then further averaged over genes and cis-regulatory regions to obtain a vector of length *L*_*r*_, representing the context-specific importance of each trans-acting regulator *i*:

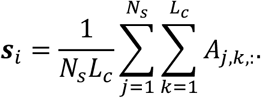

The resulting scores were ranked to prioritize trans-acting regulators for downstream gene ontology enrichment analysis.

#### Trans-regulator-TG regulatory strength

To assess the regulatory strength of trans-regulators on TGs, we focused on the 10 kbp region centered at the TSS of each candidate target gene. For a given trans-regulator k and candidate target gene j in cellular context i, the corresponding cis-and trans-regulatory sequences were input to RegFM to obtain a cross-attention matrix *A*^′^ ∈ ℝ^*L*^*c*×*Lr*, averaged across layers and heads. We then selected the column corresponding to trans-regulator k based on its index in the trans-regulatory sequence, and restricted attention to cis tokens located within the 10 kbp TSS-centered region. The regulatory score was defined by summing the attention values over these positions, yielding 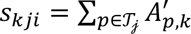, where *T*_*j*_ denotes the set of cis tokens within the TSS-centered window of gene *j*. This score quantifies the potential regulatory strength of trans-regulator *k* on gene *j* in cellular context *i*.

To split genes into target genes (TGs) and non-target genes (non-TGs), we adopted two strategies depending on data availability in each cell line. For cell lines lacking chromatin interaction data, such as H1-hESCs, we defined TGs based on ChIP-seq signal aggregated around gene promoters. Specifically, for each gene j, we considered a genomic window centered at its transcription start site (TSS) and computed a distance-weighted ChIP-seq signal by integrating all overlapping peaks within this region. The contribution of each peak was weighted by its distance to the TSS using an exponential decay function, such that closer peaks contribute more strongly. Formally, the ChIP-seq signal for gene j was defined as *S_j_* = 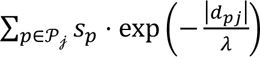, where *P_j_* denotes the set of peaks within the predefined window around gene *j*, *s*_*p*_ is the signal intensity of peak *p*, *d*_*p*j_ is the distance between the peak center and the TSS of gene *j*, and *λ* is a decay parameter controlling the contribution of distal peaks. Genes were then ranked based on *S*_j_, and the top *N* genes with the highest scores were defined as TGs, while the bottom *N* genes were defined as non-TGs. This approach prioritizes genes with strong and proximal TF binding signals, enabling a biologically informed partition of genes into likely targets and non-targets.

For cell lines paired with chromatin interaction data, we defined TGs by integrating HiChIP-derived chromatin loops with TF ChIP-seq peaks to explicitly model physical regulatory interactions. Briefly, we first identified high-confidence chromatin interactions by filtering loop calls based on ChIP enrichment, *q*-value, and normalized contact counts. Each interaction consists of two anchors, denoted as *a*_1_ and *a*_2_, with genomic coordinates (*c*, *s*, *e*). We then annotated whether each anchor overlaps TF ChIP-seq peaks and whether it is associated with gene promoters. An anchor was considered TF-bound if it overlapped at least one ChIP-seq peak. It was classified as a promoter anchor if it overlapped a promoter region defined as[*t*_j_ − *δ*, *t*_j_ + *δ*], where *t*_j_ is the TSS of gene *j* and *δ* is set to 2 kb. Formally, for an anchor a, we define a promoter indicator function *I*_prom_(*a*) = 1 if it overlaps any promoter region defined around gene TSSs |*t*_j_ − *δ*, *t*_j_ + *δ*|, and 0 otherwise.

For each loop, TGs were assigned by linking TF-bound anchors to genes whose promoters overlap either anchor of the loop. Based on these definitions, genes were classified into three categories. Promoter-TGs are genes whose promoters are directly overlapped by TF-bound anchors or connected to TF-bound promoter anchors. Enhancer-TGs are genes whose promoters are connected through chromatin interactions to TF-bound distal anchors that do not overlap promoter regions. Non-TGs are genes that are not associated with any TF-bound anchors under these criteria. This strategy integrates TF binding information with three-dimensional chromatin architecture, enabling a more precise delineation of regulatory targets and a clear separation between promoter-proximal and distal enhancer-mediated regulation.

For ChIP-seq data, we downloaded processed peak files from ENCODE, including ENCFF767MSS and ENCFF704BPD for HepG2, ENCFF196JGP, ENCFF214XPD, and ENCFF249SVT for GM12878, and ENCFF852ZRK and ENCFF148JKK for K562. These datasets were used to define TF binding signals across different cellular contexts in a unified processing framework. For HiChIP-seq data, we retrieved chromatin interaction loops at 5 kb resolution for each corresponding cell line from HiChIPdb. All loop anchors were mapped to the GRCh38 reference genome using liftOver to ensure coordinate consistency with ChIP-seq and gene annotation data. We then applied a series of quality control filters to retain high-confidence interactions. Specifically, we kept loops annotated with H3K27ac signal and further filtered them by statistical significance and interaction strength, following the criteria *q*Value < 1 × 10^−10^ and *normCount* > 5 × 10^−2^.

#### In silico knockout of trans-acting regulators

Leveraging the trans-regulatory sentence as input, RegFM naturally enables in silico knockout of specific TFs or CRs and allows quantification of their effects on genomic regions through changes in gene representations. Specifically, for each gene, we simulate the knockout of a given TF or CR by replacing its corresponding expression token with [UNK] in the trans-regulatory sentence. The original and perturbed trans-regulatory sentences, denoted as 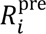 and 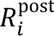, are then fed into RegFM to obtain the corresponding 768-dimensional gene embeddings *g*^pre^ and *g*^post^. The effect of the perturbation is quantified by computing the Manhattan distance between these two embeddings, which measures the extent of the embedding shift for each candidate target gene.

#### Trans-acting regulators-CRE binding strength

To quantify the binding preference of trans-acting regulators over CREs, we derived region-level attention scores from the cross-attention maps of RegFM. For each gene-centered genomic window, we extracted the attention vector corresponding to a given TF or CR from the last cross-attention layer across all cis-regulatory tokens. Let *a*_*t*,*k*_ denote the attention score assigned to cis token *t* for regulator *k*, where the attention values are averaged across heads within the last layer. Because each token corresponds to a variable-length k-mer, tokens were mapped back to genomic coordinates based on their sequence span and TSS-anchored positioning. To obtain region-level signals, we partitioned the genomic window into fixed-size bins (1 kb resolution). The binding strength of regulator *k* in bin *b* was defined as the sum of attention scores from all tokens overlapping that bin, i.e., *S*_*b*,*k*_ = ∑_*t*∈*b*_ *a*_*t*,*k*_, where *t* ∈ *b* denotes token whose genomic coordinates overlap bin *b*. This aggregation yields a continuous attention profile along the genomic region, reflecting the relative contribution of each region to the regulator.

To improve interpretability and highlight high-confidence regions, the aggregated scores were further processed using local smoothing and non-linear transformations, followed by min-max normalization to the range [0,1]. Low-intensity background signals were suppressed using a predefined cutoff, while high-attention regions were selectively amplified. The resulting normalized scores were used as a proxy for trans regulator-CRE binding strength and visualized as genomic tracks for comparison with epigenomic signals such as ChIP-seq, DNase-seq, and annotated regulatory regions.

To obtain TF binding motif scores at the k-mer level, we first ranked genomic regions based on the original k-mer-level attention scores and selected high-attention and low-attention subsets accordingly. We then performed motif scanning within these regions using HOMER, focusing on the binding profiles of specific TFs (e.g., MYC). For each region, if multiple candidate motif matches were identified, we retained the maximum motif score as the representative binding score for that region. This strategy provides a concise and robust estimate of TF binding affinity that can be directly compared with attention-derived signals.

#### Gene regulatory network inference

To infer context-specific gene regulatory networks (GRNs), a predefined set of target genes (TGs) was selected for downstream regulatory analysis. For each selected TG, RegFM was used to estimate TF-TG and TF-RE interactions based on the cross-attention module of RegFM.

Specifically, the regulatory strength between a TF and a TG was quantified by the attention weight between the gene-level representation token ([CLS]) and the corresponding TF token in the final layer of the TF-context transformer. Given a selected TG *g* and a TF *t*, the TF-TG regulatory score was defined as:

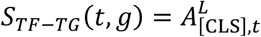

where *A*^*L*^ denotes the attention matrix from the last transformer layer when the genomic region centered around the TSS of the *TG_g_* is fed into RegFM, and 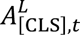 represents the attention weight from the [CLS] token to TF token *t*. To characterize TF-RE interactions, the cross-attention weights between TF tokens and genomic regulatory element tokens were extracted from the final cross-attention layer. The TF-RE interaction strength was calculated as:

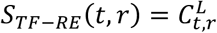

where *C*^*L*^ represents the cross-attention matrix in the final layer, and 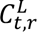 denotes the attention weight between TF *t* and regulatory element *r*. The inferred regulatory network was subsequently constructed by selecting high-confidence interactions according to predefined attention-score thresholds and edge-number constraints. Specifically, for each target gene, only the top-ranked TF-TG interactions within the specified number range were retained to generate the final GRN.

#### Motif visualization

We propose a motif visualization strategy to interpret sequence features captured by RegFM through TF-specific attention patterns. For each TF, we extract k-mer-level attention scores from the last layer and rank all tokens within gene-centered regions. High-confidence sequence signals are selected either by taking the top fraction of tokens or a fixed number of top-ranked k-mers. These tokens are mapped back to nucleotide sequences, and fixed-length subsequences are obtained by centering on each k-mer to ensure positional alignment. Based on the resulting sequence set, we construct a position weight matrix (PWM) to summarize the underlying motif pattern. Specifically, for a motif of length *L*, the PWM at position *i* for nucleotide *b* ∈ {*A*, *C*, *G*, *T*} is computed as

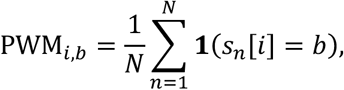

where *N* is the total number of selected sequences and *s*_*n*_[*i*] denotes the nucleotide at position *i* in the *n*-th sequence. This formulation captures the nucleotide distribution at each position and provides a probabilistic representation of attention-enriched sequence patterns. To evaluate biological consistency, the inferred PWMs are compared against known motifs in the JASPAR database^61^, and the best match is identified based on similarity. We further assess specificity by fixing the attention map derived from a given TF and comparing its PWM against motifs of other TFs. A consistently higher similarity to the matched TF indicates that the model captures TF-specific binding preferences rather than nonspecific sequence features.

## Data availability

All datasets used in this study were obtained from public sources. The pretraining transcriptomic data were downloaded from ENCODE^6^ and CELLxGENE^7^. Enhancer annotations were obtained from EnhancerAtlas^28, 29^ (http://www.singlecelldb.com/). The annotations of enhancer, silencer, insulator, and promoter elements in K562, GM12878, HeLa-S3, and HepG2 were taken from Cui et al^4^. The gene dosage sensitivity data were collected from Ni et al.^31^. The data of promoter bivalency annotation of genes were collected from Bernstein et al.^30^. Perturb-seq datasets were collected from Perturbase^36^ (http://www.perturbase.cn/), and the specific source of each dataset is provided in the Methods section. Gene embeddings from scELMo^42^ and GenePT^35^ were obtained from Zhu et al.^32^. TF binding ChIP-seq peak files used for cross-attention analysis and interpretation were collected from ENCODE. HiChIP loop data were collected from HiChIPdb^50^ (https://health.tsinghua.edu.cn/hichipdb/). Motif PWMs are collected from JASPAR database^61^. All experiment IDs from ENCODE are provided in the Supplementary Materials. All genomic regions used in this study are based on the GRCh38/hg38 reference genome or have been converted to GRCh38/hg38 using the UCSC liftOver^51^ tool. Source data are provided with this paper.

## Code availability

The source code and pretrained model of RegFM are publicly available on GitHub (ht tps://github.com/ZjGaothu/RegFM) and Hugging Face (https://huggingface.co/Deku21/Reg FM).

## Acknowledgements

This work was partly supported by the National Key Research and Development Program of China (grant nos. 2025YFC3409300, 2023YFF1204802), the National Natural Science Foundation of China (grant nos. 32550616, 62273194), Beijing Natural Science Foundation (grant no. L242026). We thank Wanwen Zeng for her early discussion in this work.

## Ethics declarations

The authors declare no competing interests.

