## Supplementary information for "RegFM: an interpretable context-aware foundation model for human transcriptional regulation"

### Supplementary Notes

#### Text S1. Implementation of genome foundation models.

We compared five genome foundation models and a sequence-based model for the gene expression prediction task: DNABERT(1), DNABERT-2(2), Nucleotide Transformer(3), Nucleic Transformer(4), EVO-2(5), and Xpresso(6). These genome foundation models provide representations of DNA sequences but do not incorporate cell type-specific information. To ensure a fair comparison, we used the officially released pretrained weights for all models and followed their tutorials for sequence-level prediction or sentence-level embedding-based inference. All models were evaluated under exactly the same setting of HVGs and cellular contexts as RegFM, ensuring a consistent and fair benchmark.

For DNA sequence inputs, short-sequence foundation models (input length < 10 kbp) were provided with 512 bp regions centered at the TSS, following the maximum input length configuration used in DNABERT. For Xpresso, which is a convolution-based model, we used its original input setting, i.e., 7 kbp upstream and 3.5 kbp downstream of the TSS.

For DNABERT, we evaluated all four pretrained variants released in the original work. As shown in Fig. S4, different k-mer settings showed no significant differences in gene expression modeling performance. We therefore report results using the 6-mer model in the main text. For the Nucleotide Transformer, we downloaded the nucleotide-transformer-500m-human-ref model from Hugging Face (https://huggingface.co/InstaDeepAI/nucleotide-transformer-500m-human-ref) and used it as the pretrained checkpoint for fine-tuning. For EVO-2, which is based on the HyenaDNA architecture and supports long-context modeling, we used the EVO-2 7B model from Hugging Face (https://huggingface.co/arcinstitute/evo2_7b). We evaluated three input sequence lengths centered at the TSS: 8,192 bp, 200 kbp, and 650 kbp. Sequence-level embeddings were obtained via mean pooling and used to predict gene expression. During fine-tuning, we froze the EVO-2 backbone and trained only a new prediction head on top of the DNA region embeddings. As shown in Fig. S5a-b, the model with 8,192 bp input achieved better performance than the long-sequence models. This suggests that mean pooling may over-smooth representations over long contexts, potentially diluting cis-regulatory signals around the TSS.

Furthermore, we concatenated EVO-2 DNA embeddings with normalized TF/CR embeddings to match the input information used in RegFM. As shown in Fig. S5c-e, this consistently improved performance across all three sequence lengths. However, the results still lagged significantly behind RegFM, suggesting that simply fine-tuning a prediction head on engineered features is insufficient to capture how TFs and cis-regulatory elements interact with DNA sequence to regulate gene expression.

#### Text S2. Implementation of CREATE in CRE prediction task.

In the CRE prediction task, in addition to genome foundation models based on DNA sequences and classical machine learning models using k-mer features, we further compared against state-of-the-art CRE-specific methods, including CREATE and CREATE-seq.

Specifically, for the EnhancerAtlas(7) dataset, we did not include CREATE because its required openness scores cannot be consistently derived following the original implementation. However, in the insulator identification task, openness scores are available, and the original CREATE paper reports that CREATE (seq+open) performs very similarly to the full CREATE model. We therefore evaluate CREATE (seq+open), denoted as CREATE, alongside CREATE-seq, which uses DNA sequence only.

All evaluations were conducted on the hg38 reference genome. We used LiftOver(8) to map CRE regions to hg38 coordinates. Openness scores were retained from the original positions in OpenAnnotate(9, 10) and aligned to hg38 at base-pair resolution to match the corresponding DNA sequences. As shown in Fig. S11, CREATE consistently outperforms CREATE-seq, benefiting from epigenomic signals that provide inherent cell type awareness. This advantage is also reflected in Fig. 3d, where CREATE shows substantially stronger performance than other baseline methods.

#### Text S3. Strategies of RegFM for predicting transcriptional responses to genetic perturbations.

To leverage the context-specific gene embeddings generated by RegFM for predicting transcriptional responses to genetic perturbations, we integrated RegFM embeddings into the Scouter framework using two strategies as illustrated in Fig. S2: (i) replacing the perturbation embedding, where the original perturbation embedding was directly replaced with the RegFM-derived gene embedding under the corresponding cellular context; and (ii) attention-based fusion, where the static gene embedding provided by Scouter was fused with the RegFM-derived gene embedding through an adaptive fusion module. Specifically, given the Scouter embedding $e_s$ and RegFM embedding $e_c$, we first concatenated them and computed adaptive fusion weights using an attention network:

$$\alpha = \mathrm{Softmax}\big(f([e_s; e_c])\big)$$

The two embeddings were then projected into a shared latent space and combined by a weighted sum:

$$e_f = \alpha_1 W_s e_s + \alpha_2 W_c e_c$$

where $e_f$ represents the final fused gene embedding used for downstream perturbation response prediction. Based on these two strategies, we were able to evaluate the effectiveness of RegFM embeddings through strategy (i) (Fig. 4c-d), and further demonstrate that RegFM embeddings, which capture transcriptional regulatory knowledge learned from omics data, provide complementary information to GenePT gene embeddings derived from textual knowledge through strategy (ii) (Fig. 4e-g).

#### Text S4. Details of cross cellular context prediction of transcriptional perturbation responses of RegFM.

For cross-dataset or cross-condition prediction, a leave-one-out strategy was adopted to evaluate the generalization ability of RegFM to unseen cellular contexts. Specifically, the five Perturb-seq datasets were integrated as follows: (1) using Scanpy(11), genes were aligned across datasets by selecting the intersection of shared genes as anchors; (2) batch effects were corrected using Harmony(12); and (3) one dataset was held out as the test set, while the remaining four datasets were integrated and used for model training.

Model training and prediction were performed within the Scouter framework. During each training iteration, the unperturbed cells from each training dataset were used as dataset-specific control references to model perturbation-induced transcriptional responses under different cellular contexts. For evaluation, the unperturbed cells from the held-out dataset were used as controls, and the trained model was applied to predict transcriptional responses to perturbations in this unseen context. This design evaluates the ability of RegFM to generalize beyond observed perturbation–context combinations and predict gene expression responses in previously unseen cellular environments. All other procedures were kept consistent with the single-dataset prediction setting.

### Supplementary Figures

#### Fig. S1

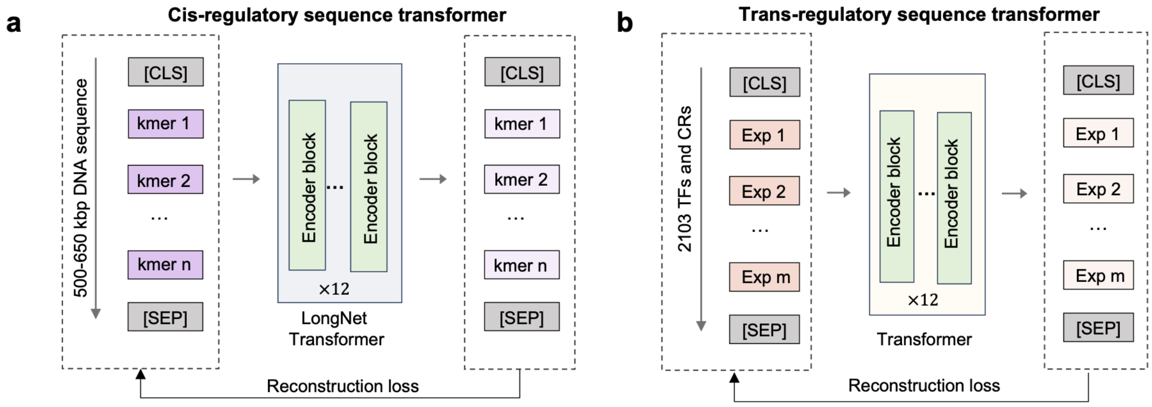

**Fig S1. Self-supervised pretraining strategy for cis-transformer (Cis-T) and trans-transformer (Trans-T). a**, The DNA transformer module takes long-range DNA regions exceeding 500 kbp as input and adopts a LongNet-based transformer encoder block for masked language modeling (MLM) pretraining. **b,** The regulator transformer module takes TF/CR sequences as input, where each token corresponds to a binned discretized expression value of a TF/CR.

#### Fig. S2

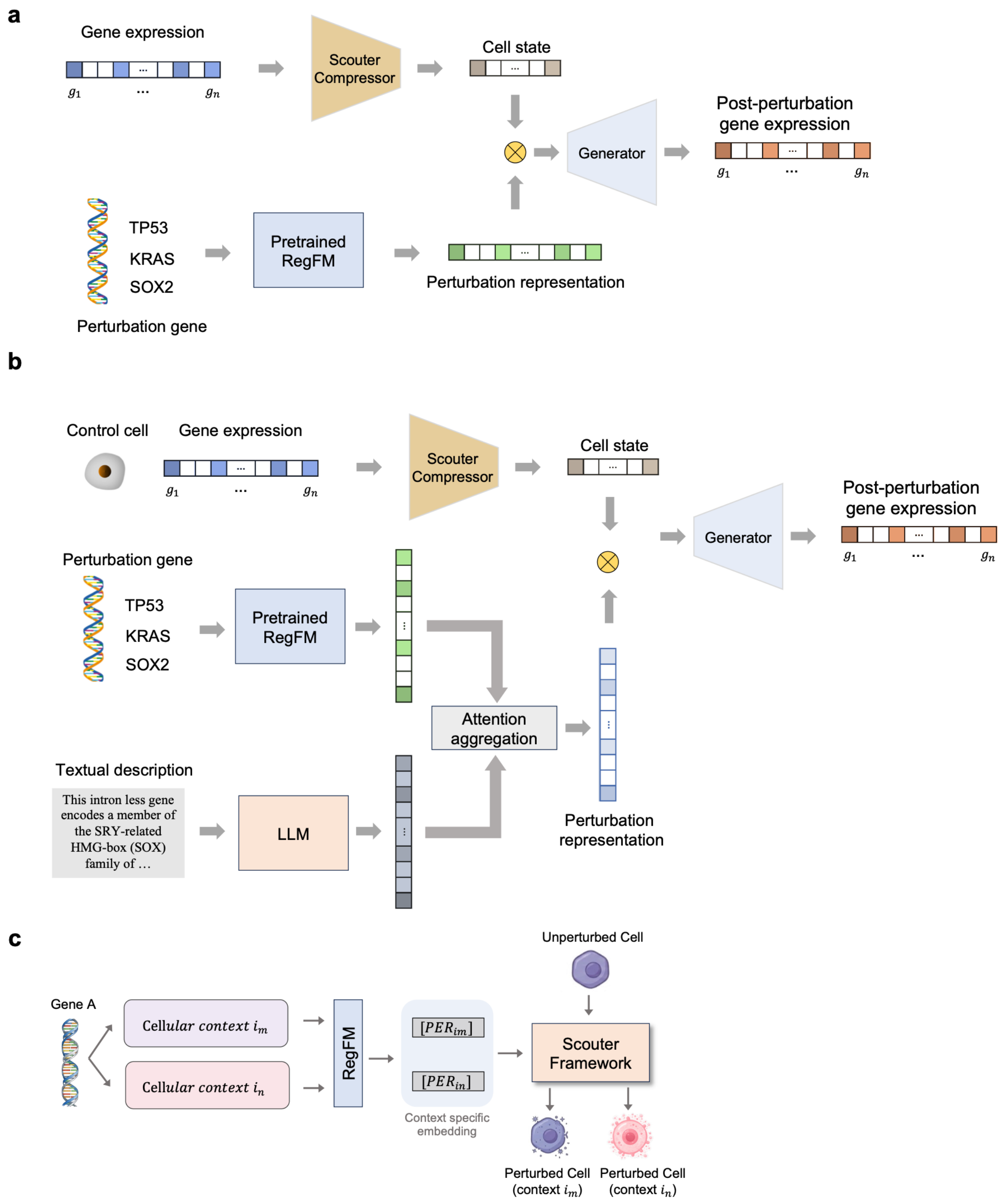

**Fig. S2. Two fine-tuning strategies of RegFM for predicting transcriptional responses to genetic perturbations.** a, RegFM generates perturbation gene embeddings under the corresponding cellular context. The perturbation gene embeddings are concatenated with cell embeddings generated by the Scouter(13) compressor and passed to a generator to predict post-perturbation gene expression. b, RegFM perturbation gene embeddings are integrated with Scouter-provided gene embeddings through an attention-based fusion module, followed by a prediction framework similar to that in (a). c, Workflow for cross-cellular context prediction of post-perturbation states using RegFM.

### Fig. S3

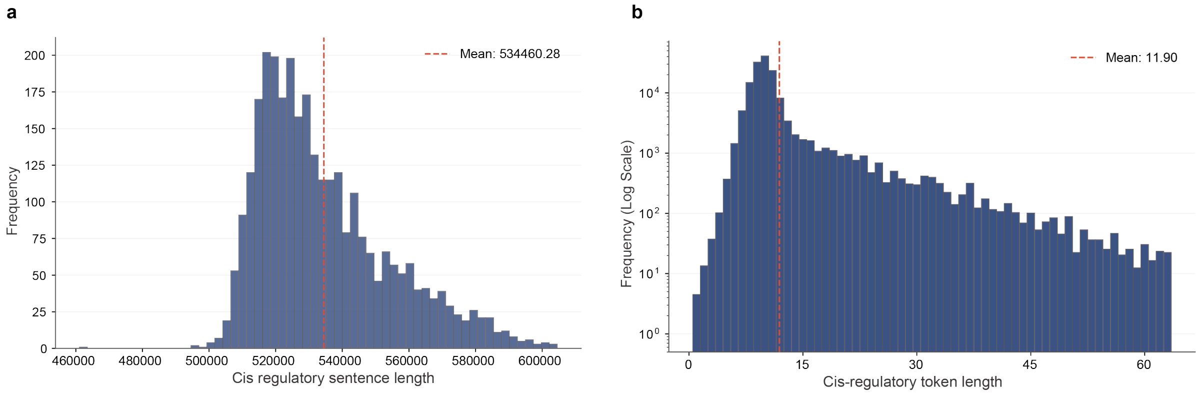

**Fig. S3. Distribution of DNA sequence lengths and token lengths of the BPE tokenizer.** a, Histogram of genomic lengths covered by 71,680 Cis-T module tokens after tokenizing the 325-kbp regions surrounding the TSSs of 3,000 HVGs, with a mean coverage of approximately 524 kbp. b, Histogram of k-mer lengths in the Cis-T tokenizer vocabulary, with a mean length of 11.9 bp.

### Fig. S4

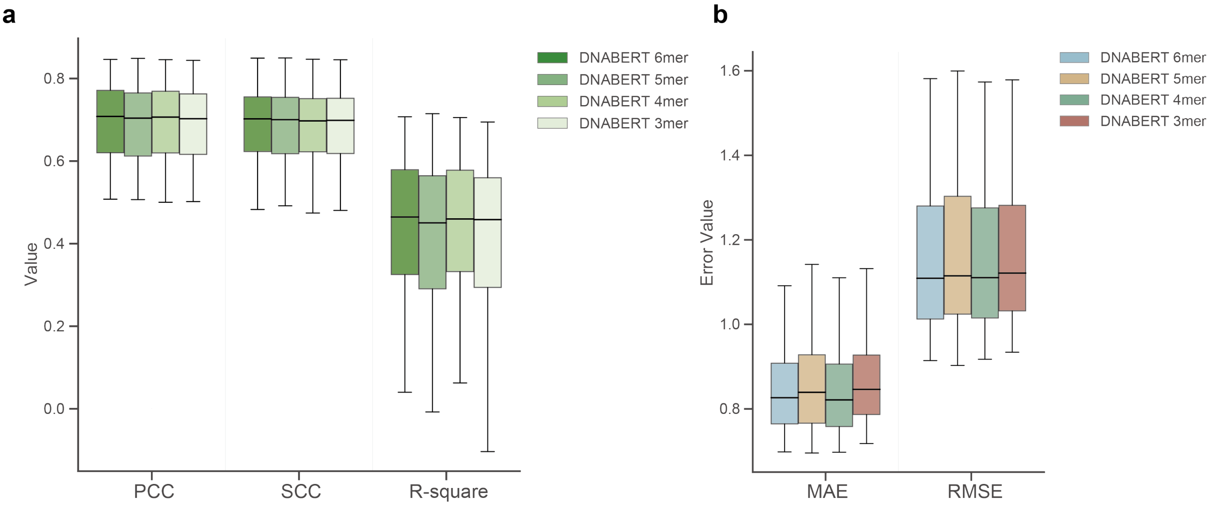

**Fig. S4. Performance of pretrained DNABERT(1) with different k-mer settings for predicting HVG expression.** a, Box plots of prediction performance (PCC, SCC, and R-square) for DNABERT (3-mer, 4-mer, 5-mer, and 6-mer) across 3,000 HVGs in 56 unseen cellular contexts. b, Box plots of prediction errors (MAE and RMSE) for DNABERT (3-mer, 4-mer, 5-mer, and 6-mer) for each of the 3,000 HVGs across 56 unseen cellular contexts.

#### Fig. S5

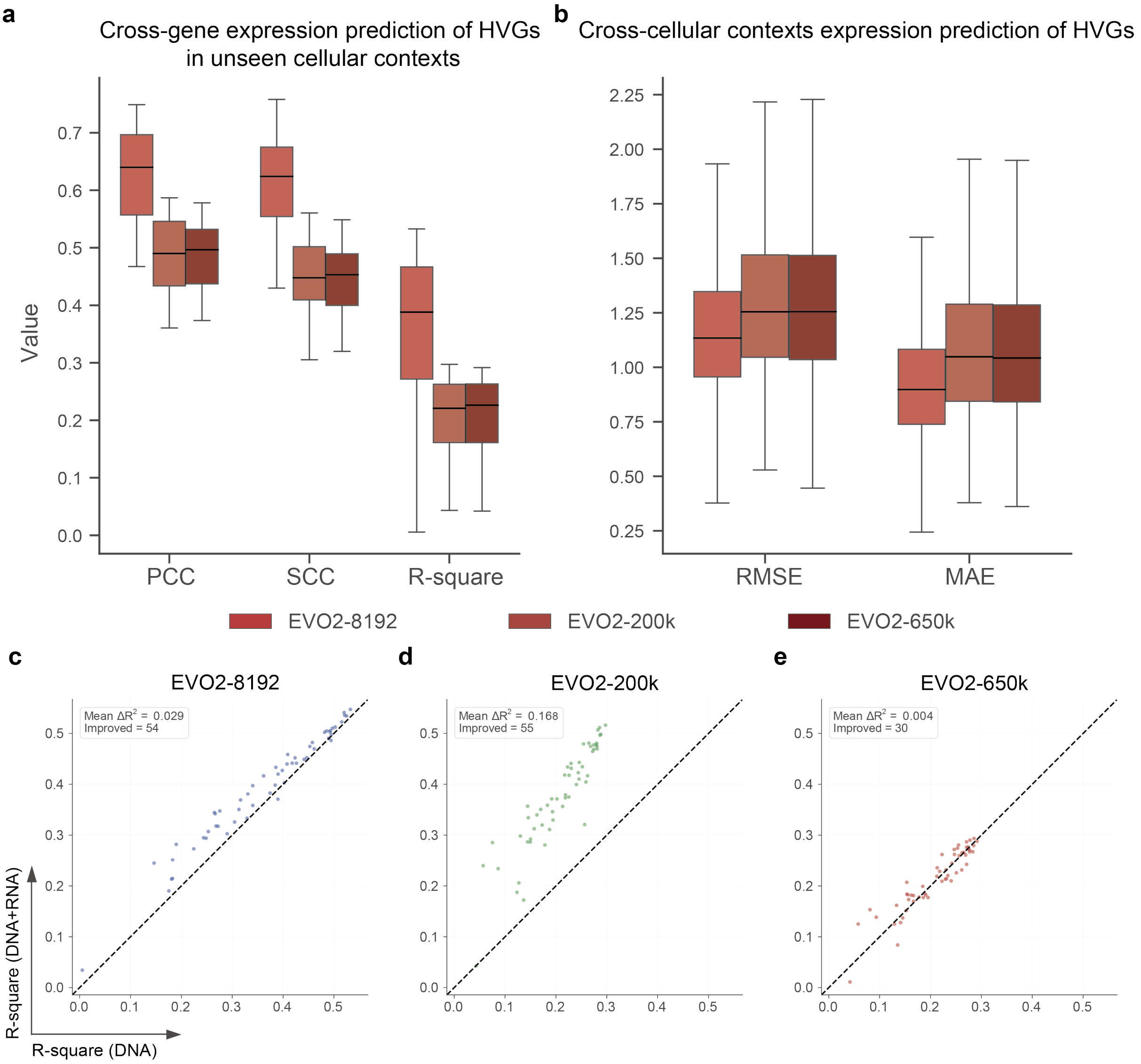

**Fig. S5. Performance of pretrained EVO-2(5) with different DNA input sequence lengths for predicting HVG expression.** a, Box plots of prediction performance (PCC, SCC, and R-square) for EVO-2 using sequence lengths of 8,192 bp, 200 kbp, and 650 kbp around the TSS across 3,000 HVGs in 56 unseen cellular contexts. b, Box plots of prediction errors (MAE and RMSE) for EVO-2 using sequence lengths of 8,192 bp, 200 kbp, and 650 kbp around the TSS for each of the 3,000 HVGs across 56 unseen cellular contexts. c–e, Scatter plots comparing the prediction performance (R-square) of EVO-2 using DNA sequence only (x-axis) and EVO-2 using the same 2,103 TF/CR expression features as input for predicting 3,000 HVGs across 56 unseen cellular contexts: (c) 8,192 bp, (d) 200 kbp, and (e) 650 kbp.

#### Fig. S6

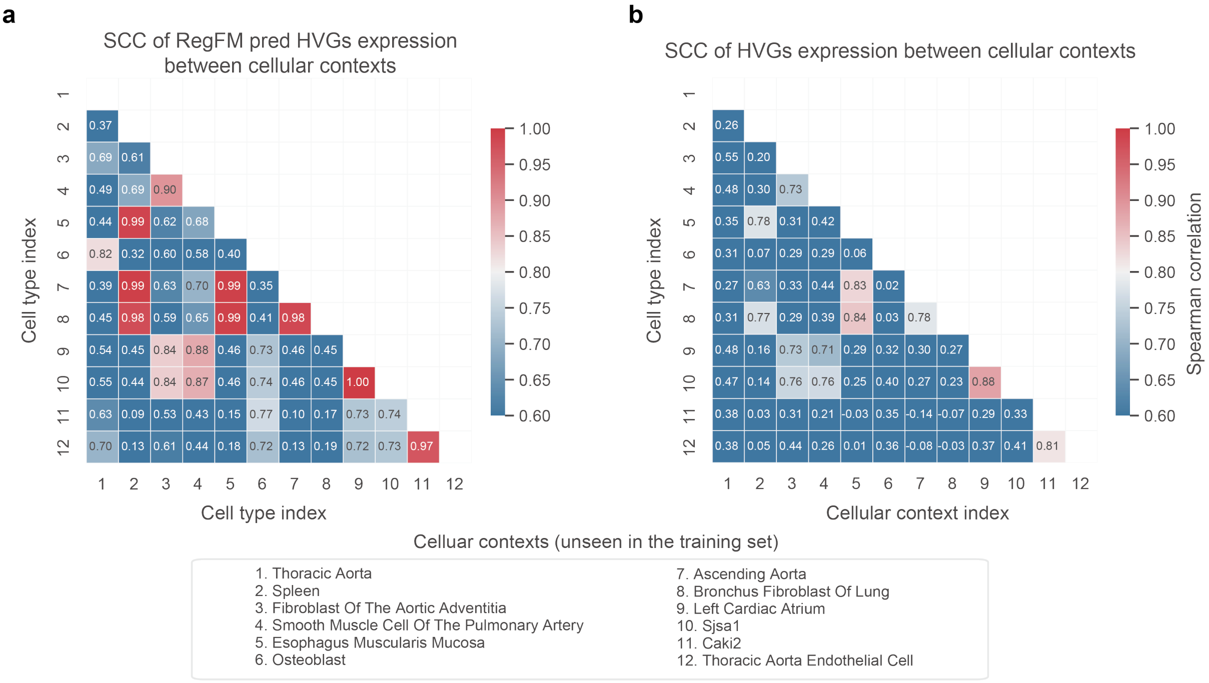

**Fig. S6. Cell type–specific gene expression patterns learned by RegFM.** a, Heatmap showing the similarity (SCC) of RegFM-predicted expression profiles for 3,000 HVGs across 12 randomly selected unseen cellular contexts. b, Heatmap showing the similarity (SCC) of the ground-truth expression profiles (normalized TPM values) for the same 3,000 HVGs across the same 12 unseen cellular contexts.

#### Fig. S7

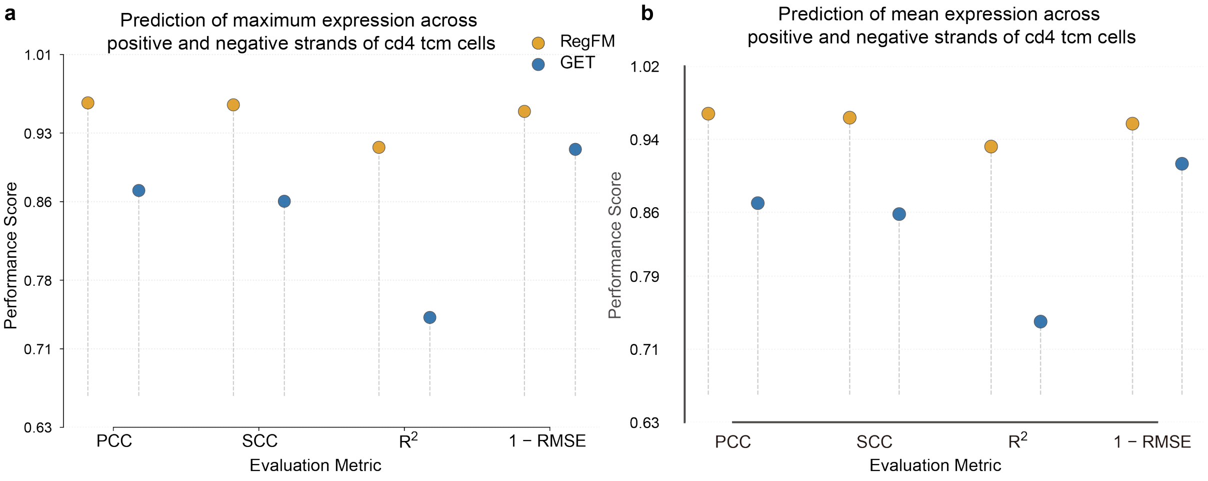

**Fig. S7. Comparison of RegFM and GET(14) for predicting the (a) maximum and (b) mean expression values between positive and negative strands in CD4 Tcm cells (PCC, SCC, $R^2$, and 1−RMSE).**

### Fig. S8

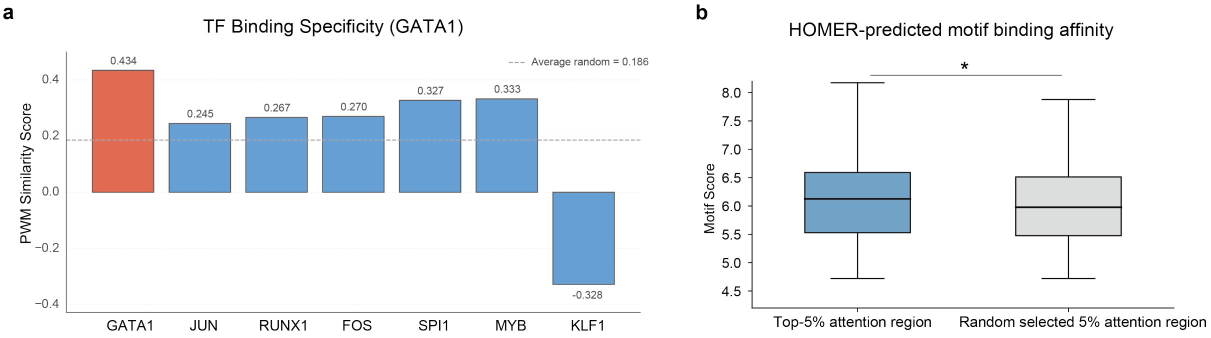

**Fig. S8. Interpretation of the cross-attention score between TF/CR and CREs.** a, Bar plot of PWM similarity between the GATA1 PWM inferred from high-attention regions of RegFM and known TF motifs from the JASPAR database(15). The x-axis shows the similarity between the inferred PWM and PWMs of different known TF motifs. b, Box plot of Homer scanned motif scores comparing top 1% and bottom 1% attention-ranked regions around the TSS of BAK1 in K562 cell line.

### Fig. S9

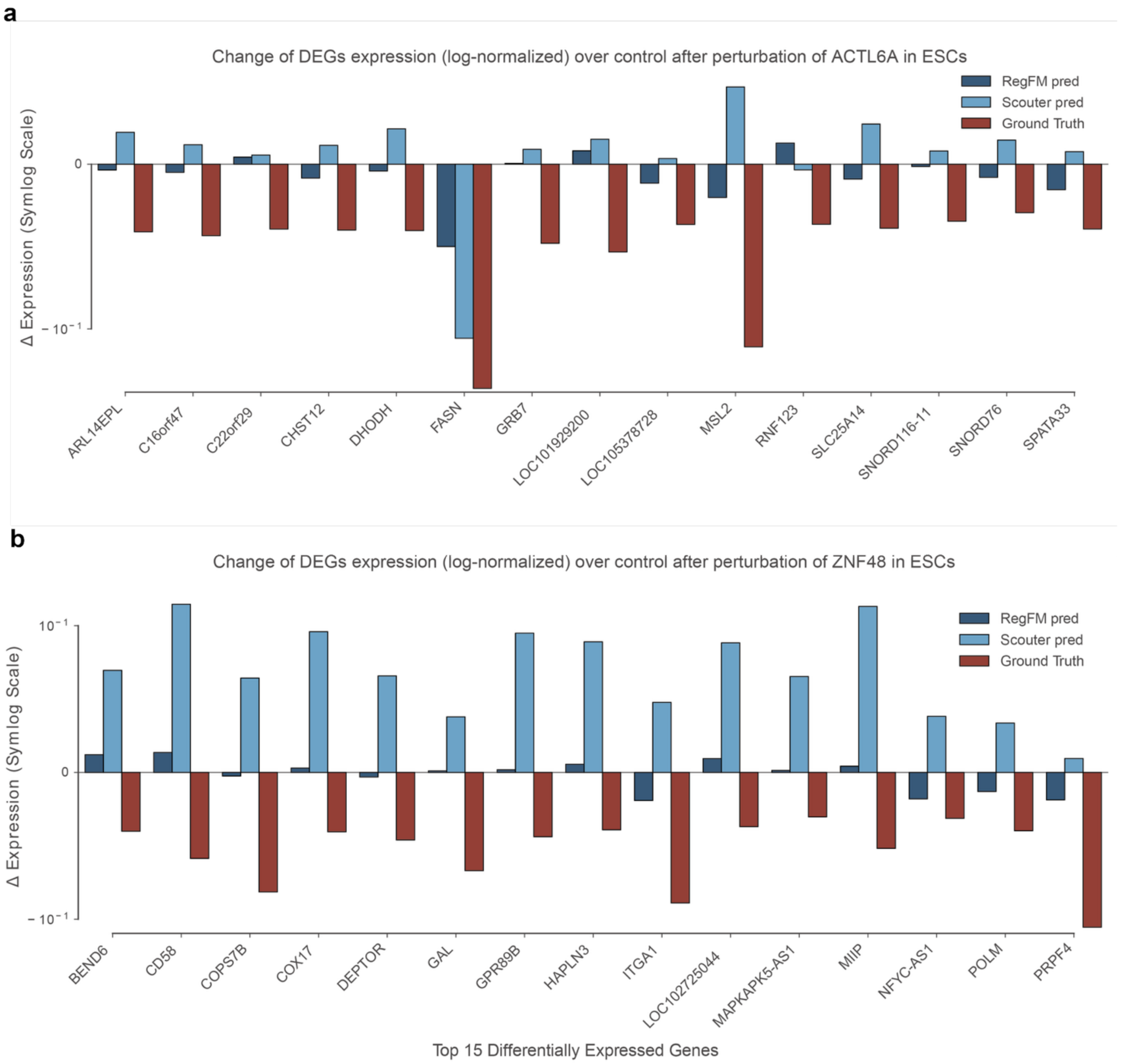

**Fig. S9. Fine-tuning RegFM for predicting transcriptional responses to genetic perturbations.** a, Log-transformed predicted expression changes of fine-tuned RegFM, Scouter(13), and observed expression changes for the top 15 DEGs in ESCs following ACTL6A perturbation. b, Log-transformed predicted expression changes of fine-tuned RegFM, Scouter, and observed expression changes for the top 15 DEGs in ESCs following ZNF48 perturbation.

### Fig. S10

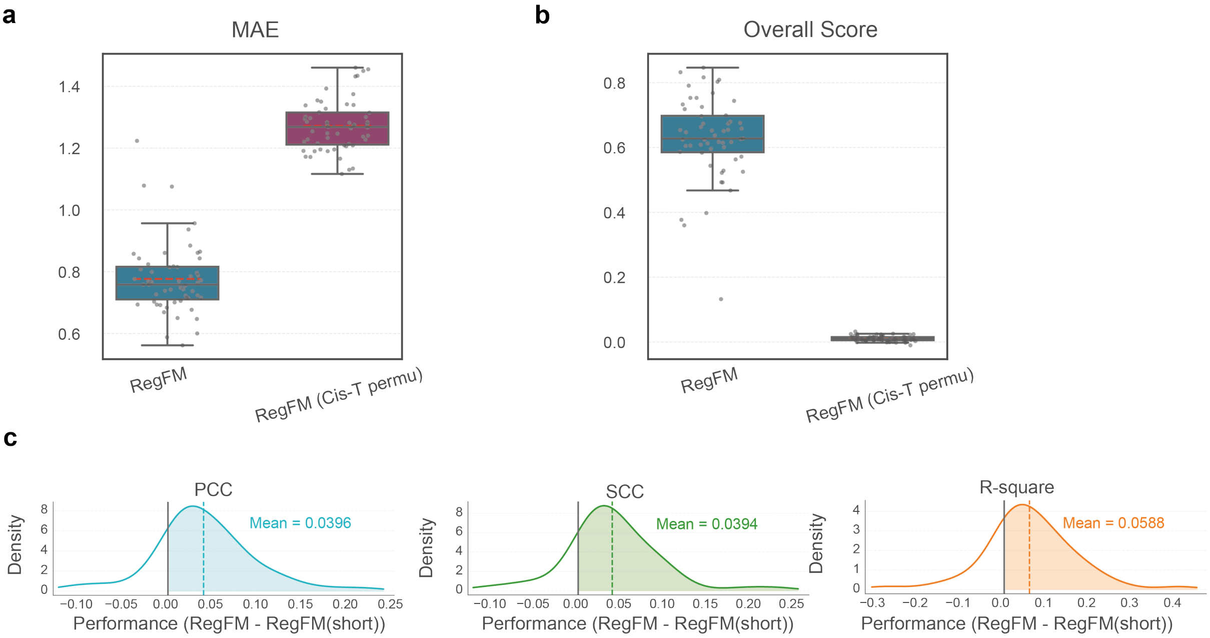

**Fig. S10. Ablation study of ReFM.** a-b, Permutation ablation analysis of the Cis-T module in RegFM**.** Performance of RegFM before and after Cis-T permutation for predicting expression of 3,000 HVGs across 56 unseen cellular contexts: (a) MAE, (b) overall score $(0.25 \times (PCC + SCC) + 0.5 \times R^2)$. c, Distribution of gene expression prediction performance across 56 unseen cell lines and tissues for RegFM compared with RegFM using short DNA sequence inputs.

#### Fig. S11

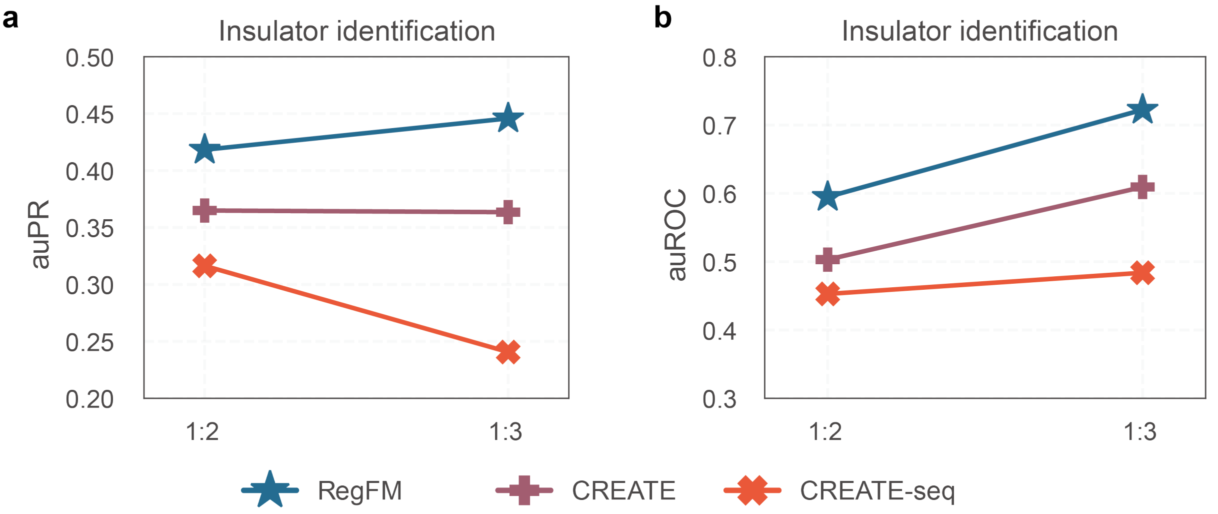

**Fig. S11. Comparison of RegFM and CREATE(16) for insulator identification across HepG2, K562, GM12878, and HeLa-S3.** a, auPR of RegFM, CREATE, and CREATE-seq for insulator identification under different positive-to-negative sample ratios (1:2 and 1:3). b, auROC of RegFM, CREATE, and CREATE-seq for insulator identification under different positive-to-negative sample ratios (1:2 and 1:3).

#### Fig. S12

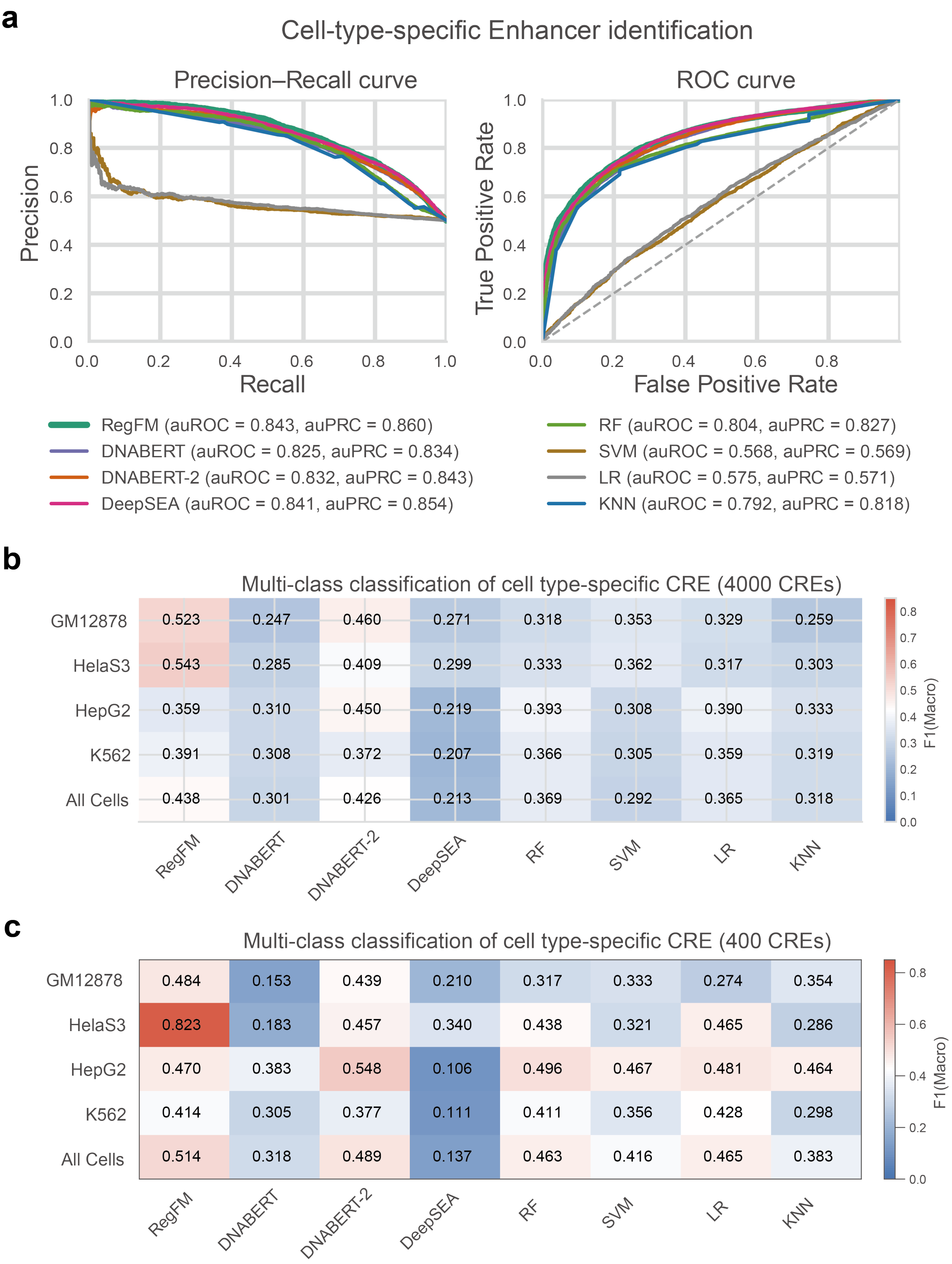

**Fig. S12. Performance of fine-tuned RegFM and baseline methods for CRE identification.** a, ROC and precision–recall curves for cell-type-specific enhancer identification. b, Macro-averaged F1 score of RegFM and baseline methods for cell-type-specific multi-class classification of regulatory elements, including enhancers, silencers, insulators, and background regions (4,000 CREs per cellular context). c, Macro-averaged F1 score of RegFM and baseline methods for cell-type-specific multi-class classification of regulatory elements, including enhancers, silencers, insulators, and background regions (400 CREs per cellular context).

#### Supplementary Tables

Table S1. List of cell lines/tissues from ENCODE.

Table S2. List of cell types from CELLxGENE.

Table S3. Experiment accessions for bulk RNA-seq data used in the ENCODE project.

Table S4. List of TF/CRs used for RegFM training.

Table S5. Datasets used for genetic perturbation response prediction experiments.

Table S6. List of the 44 cell lines and tissues with enhancer annotations collected from EnhancerAtlas.
